# Spike antibodies targeting GRP78 predispose to cardiovascular complications compared to Dengue

**DOI:** 10.64898/2026.05.20.726568

**Authors:** Supratim Sarker, Trina Roy, Abinash Mallick, Sayantan Das, Sarakam Dharma Teja, Arun Bandyopadhyay, Abhishek De, Surajit Gorai, Subhajit Biswas

**Author notes:** Joint corresponding authors; Lead contact: Subhajit Biswas. Supratim Sarker and Trina Roy contributed equally. Author, (S.S.), (T.R.), (A.M.), (S.D.), (A.B.) (A.D.) (S.D.T.) (S.G.). Materials Availability: This study did not generate new unique reagents. Data and Code Availability: **The data generated from mass spectrometric analyses of serum sample from healthy controls and affected individuals diagnosed with COVID-19 or Dengue, have been deposited in PRIDE repository with dataset identifier PXD072757 and 10.6019/PXD072757**^1^ **under the Project Name**: Comparative plasma proteomics of COVID-19 and Dengue affected individuals with relevance to cardiovascular-associated **pathways;** using ProteomeXchange^2^. Link: https://proteomecentral.proteomexchange.org/ui?pxid=PXD072757.

## Abstract

One major aftermath of COVID-19 pandemic is cardiovascular consequences. SARS-CoV-2 binds to ACE2 and downregulates vasodilation. Dengue favors hypotension by weakening endothelial glycocalyx leading to plasma leakage. C1q levels, immune complexes (ICs), and proteomic profiles in serum samples from 52 COVID-19 and 19 pre-pandemic Dengue cases were studied. Unlike Dengue, COVID-19 serums showed elevated coagulation proteins promoting vaso-occlusion and peripheral artery diseases.

The stress-induced chaperone and atherosclerosis marker, GRP78 (gene/ protein) was found upregulated upon SARS-CoV-2 spike expression in cardiac/ lung cell lines. Elevated GRP78 levels were also observed in serum samples from COVID-19-diagnosed individuals and subjects with myocardial infarction (MI) in post COVID-era.

Surprisingly, spike antibodies (Abs) showed cross-binding to GRP78 and possibly contributed to the observed higher-level ICs in COVID-19 serums (cardiovascular embolism?). Co-localization studies showed that spike Abs (analogous to pro-atherosclerotic GRP78 auto-Abs) could directly bind to upregulated cellular GRP78 (type II hypersensitivity?). Both pathways could worsen vascular injury and atherosclerosis, leading to cardiac complications in COVID-19 cases with narrowed vessels.

## 1. INTRODUCTION

The year 2019 marked the incidence of a wave of pneumonia cases caused by a novel coronavirus SARS-CoV-2 in the Wuhan district of Hubei Province, China, ^3^ that soon escalated to the global COVID-19 pandemic with over 700 million cases and 7 million deaths reported as on April, 2024^4^. There has been sufficient evidence revealing that many survivors of this pandemic had been suffering from a wide range of persisting health issues, known as post-COVID syndrome or “Long COVID”, following their initial recovery^5, 6^. Various mechanisms including persisting infection, interferon (IFN) response, thrombo-inflammation, and autoimmune processes had been implicated as the underlying pathophysiology of Long COVID but a clear picture is still to emerge^5,7,8,9^.

One of the common symptom manifestations associated with COVID-19 had been cardiac complications^6,10^. Myocardial injury, myocarditis, acute heart failure, and cardiomyopathy had all been reported during the acute phase of COVID-19^11^. Besides clinical/subclinical myocardial damages, SARS-CoV-2 infection also caused myocarditis with severe left ventricular diastolic dysfunction in a larger number of cases^12,13^. Many young, healthy individuals who had COVID-19 were subsequently found to have unsuspected cardiac pathology^14^.

Thrombotic coagulation problems had also been frequently encountered, however exact reasons for such coagulopathy in COVID-19 are still being elucidated^15^. For instance, acute inflammation caused by SARS-CoV-2 manifested as endothelial dysfunction, platelet activation, and hypercoagulability^16^. Diffuse alveolar and pulmonary interstitial inflammation in COVID-19 had been shown to be caused by an immunological mechanism that involved a “macrophage activation syndrome”-like condition which led to widespread immune thrombosis^17^.

According to several studies, SARS-CoV-2 infection disrupted the balance between ACE/ACE2 and Angiotensin (AT)-II/AT-1-7, which could affect blood pressure (BP) and accelerate the development of arterial hypertension by vasoconstriction and down-regulated vasodilation pathways^18,19^. Thus, compared to individuals without hypertension, COVID-19 cases with hypertension demonstrated worse disease progression and higher disease severity^20^. Study indicated that prolonged COVID-19 had an impact on the clinical progression of conditions linked to hypertension, including cardiovascular diseases, kidney diseases, and endocrine problems^21^.

The genome of SARS-CoV-2 encodes four structural proteins: S (spike), E (envelope), M (membrane) and N (nucleocapsid), all playing essential roles in different phases of viral infection. Among the structural proteins, the spike majorly accounts for viral interaction with host angiotensin-converting enzyme 2 (ACE2) and subsequent virus entry^22^. Besides acting as a viral receptor, ACE2 also functions as an antagonist to ACE1 (Angiotensin-converting enzyme) of the Renin-Angiotensin-Aldosterone System (RAAS)^23^, which regulates BP. Proteolytic cleavage of ACE2, on binding with spike protein leads to fragile and inferior vascular conditions, thus predisposing to cardiovascular complications. Deficiency of ACE2 coupled with upsurge of fibrinogen/ fibrin breakdown products^24^, and D-dimer formation^25^, heightened the risk of COVID-19 affected individuals with acute respiratory disease syndrome (ARDS), to encounter thromboembolic events^26^.

Unlike the anti-viral protein (spike-specific IgG) that drives the acute disease severity^27^, Long COVID is hypothesized to be additionally maneuvered by the presence of self-antigen targeting autoantibodies (auto-Abs against components of the immune system, cardiovascular system; thyroid auto-Abs and yet unidentified other auto-Abs)^28^. Besides the consequences of direct binding of spike to ACE2 receptor, there had been both clinical and experimental data suggesting that the presence of autoimmune Abs can also intensify the risk of cardiovascular disease pathogenesis^29^ and other adverse effects via immune complex (IC) formation^30,31^ . However, the origin and role of various auto-Abs in Long COVID pathophysiology remains an elusive area of active investigation^32^.

In this regard, it is noteworthy that certain studies have suggested that accumulation of ICs in different areas of the heart had been pivotal in autoimmune-associated inflammatory reactions and activation of body’s complement system^33^. Chronic inflammation characterized by accumulation of blood monocytes; up-regulation of intercellular adhesion molecules; release of pro-inflammatory cytokines; and production of matrix degrading enzymes, acted as essential contributing factors towards vascular endothelial dysfunction, critical in the early onset of cardiac disorders^34^.

IC-related illnesses are known to make affected individuals more vulnerable to platelet-mediated thrombotic events. So, it can be hypothesized that IC formation and subsequent related events have the potential to cause/augment intravascular coagulation pathology in case of SARS-CoV-2 infection. While Dengue virus (DV) and SARS-CoV-2 enter the body through different routes, both infections cause systemic illnesses and share some common clinical manifestations, including fever, headache, myalgia, and gastrointestinal symptoms. While bleeding tendencies are typically linked to Dengue, the development of micro- and macro-thrombi had been a more predominant sign of severe COVID-19^35^.

In this study, we have proposed two possible mechanisms that might be contributing towards the underlying pathophysiology of the cardiovascular manifestations of Long COVID. For this, we have implemented a combination of serum analysis, mass spectrometric studies and cell culture models.

## 2. MATERIALS AND METHODS

### Study subjects

Fifty-two archived serum samples from clinically and laboratory-confirmed (swab reverse transcription-polymerase chain reaction [RT-PCR]-positive) COVID-19 cases were collected from Apollo Multispecialty Hospital, Kolkata, from September 2020 to August 2022. All affected individuals showed mild to severe COVID-19 symptoms but were discharged from the hospital eventually on recovery. Similarly, archived serum samples from 19 Pre COVID-19 pandemic Dengue cases were collected from Apollo Multispecialty Hospital in 2017 (Supplementary table 1). Fourteen archived serum samples from affected individuals with myocardial infarction (MI) were collected from Calcutta National Medical College (CNMC) in October 2023 (Supplementary table 2). Eleven archived serum samples were obtained from apparently healthy individuals from Apollo Multispecialty Hospital, Kolkata in March, 2020. The study was approved by the respective Institutional Ethical Committees of the previously mentioned hospitals and the Council of Scientific & Industrial Research-Indian Institute of Chemical Biology (CSIR-IICB), Kolkata. Written informed consent (in their native language) was obtained from all individual participants included in this study. All experiments were carried out as per relevant guidelines and regulations.

### ELISA

Human C1q solid-phase sandwich ELISA kit (sensitivity 0.08 ng/ml), purchased from Thermo Fisher Scientific (Cat no: #**BMS2099)** was used to quantify Human C1q. Circulating IC (CIC)-C1q level in affected and healthy serum samples was quantified by CIC C1q ELISA kit (purchased from DRG International, Cat no: **EIA-3169**) having a threshold value of 16 μg Eq/ml. Standards, control and samples were added to the pre-coated microplate and following steps were followed as per manufacturer’s instructions. On addition of the HRP – labeled anti-human IgG (detector) antibody (Ab), a sandwich is created between the detector and the antigen-Ab (C1q-anti C1q or CIC-C1q complex respectively). TMB substrate reacts with the enzyme-Ab-target complex, to generate a colored product (wavelength measured at 450 nm) that directly relates to the concentration of the target. The concentration of target in the samples was then determined from the derived standard curve. All the samples were run in duplicate and analyzed individually.

### Cell culture and transfection

H9c2, A549, and HEK293T cell lines were obtained from the National Centre for Cell Science, India. All these cell lines were cultured in DMEM media (Sigma) supplemented with 10% FBS (Gibco), Pen-Strep and L-glutamine mix (Sigma), and amphotericin B at 2.5 ug/ml (Gibco). These cells were cultured at 37[ at 5% CO2/95% air. During sub-culturing, cells were washed with PBS (1x) and dissociated from the substratum using trypsin-ethylene di-amine tetra-acetic acid (EDTA) (1x) (Gibco). Transfection experiments with pUNO 1 spike plasmid (Invivogen) were done using Fugene transfection reagent as per the manufacture’s protocol.

### Mass spectrometry

#### In Solution Sample Preparation

Proteins from healthy control and affected individual serum samples were precipitated using 10% trichloroacetic acid (TCA; Sigma, #T9159) in acetone (Sigma, #650501) at a 1:9 (v/v) ratio. Samples were incubated overnight at −20 °C and centrifuged at 15,000 × g, 4 °C for 15 min. The resulting protein pellet was resuspended in 1% sodium deoxycholate (Sigma) prepared in 50 mM ammonium bicarbonate (Sigma). Protein concentration was determined, and 30 μg of protein was used for downstream processing. Samples were reduced in 20 mM freshly prepared dithiothreitol (DTT; Sigma, #10197777001) by incubating at 56 °C for 40 min. Samples were alkylated using 50 mM freshly prepared iodoacetamide (IAA; Sigma, #I6125), followed by incubation at room temperature for 40 min in the dark. Proteins were digested overnight at 37 °C with 1 μg trypsin protease (Pierce Thermo, #90057) under mild agitation.

#### In-Gel Sample Preparation

For in-gel digestion, protein bands of interest were excised from Coomassie Brilliant Blue G-250 – stained polyacrylamide gels. Post destaining, the gel pieces were reduction and alkylation using DTT and IAA as described above; followed by dehydration in 100% acetonitrile. The air-dried gel pieces were then rehydrated in 25 mM ammonium bicarbonate containing 1 μg trypsin protease and incubated overnight at 37 °C for digestion.

#### Mass-spectra Generation and Analyses

Lyophilized peptide samples were freshly reconstituted in 0.1% Formic Acid and analyzed by tandem mass spectrometry (MS/MS) on an Orbitrap mass spectrometer (LTQ-XL, Thermo Fisher Scientific) coupled to an Easy-nLC 1000 system using a C18 Easy-Spray nano column (3 μm particle size). The peptides were separated over a 145-min water and acetonitrile gradient prior to ionization and high-resolution mass analysis for protein identification and label-free quantification.

Raw files were processed using Proteome Discoverer v1.4 (Thermo Scientific) and searched against protein database using Mascot. Comparative quantitative analysis across experimental groups was performed using SIEVE (Thermo Fisher Scientific), and differentially expressed proteins were subjected to GO and pathway enrichment analyses using PANTHER (http://www.pantherdb.org/)^36^, and STRING databases (https://string-db.org)^38^.

#### Immunofluorescence and confocal microscopy

H9c2, A549, and HEK293T cells were cultured on sterile coverslips and fixed with 4% paraformaldehyde (PFA) in 1× PBS for 10 minutes at room temperature (RT). Permeabilization was carried out using 1× PBST (0.1% Tween-20 in 1× PBS) for 15 minutes at RT. Subsequently, cells were blocked in 1% bovine serum albumin (BSA) prepared in PBST for 1 hour at RT on a rocking platform to reduce non-specific binding.

After blocking, cells were incubated for 1 hour at RT with primary antibodies diluted in 1% BSA in PBST. The following Ab dilutions were used: anti-Spike S1 (1:50, Abcam, ab275759) and anti-GRP78 (1:400, Abcam, ab21685). For co-localization studies, Spike S1 Ab was conjugated with Alexa Fluor 488 (Invitrogen, A20181), and GRP78 Ab was conjugated with Alexa Fluor 568 (Invitrogen, A10238).

After primary Ab incubation, cells were washed three times with PBST (5-minute intervals per wash). For standard immunofluorescence (excluding co-localization), cells were further incubated in the dark for 30 minutes at RT with an Alexa Fluor 488-conjugated secondary Ab (Cell Signaling Technology, 4418), diluted 1:1000 in 1% BSA in PBST (secondary Ab was not used for colocalization study). Following secondary incubation, cells were washed four times with PBST at 5-minute intervals on a rocking platform.

Following final washes, coverslips were air-dried and mounted onto glass slides using ProLong™ Diamond Antifade Mountant with DAPI (Invitrogen) for nuclear staining. Images were acquired using a Leica TCS SP8 confocal microscope equipped with an oil immersion objective lens. Image acquisition and processing were performed using LAS X software.

#### Immuno-blotting

H9c2, A549, and HEK293T cells were transfected with the pUNO1-Spike plasmid and collected for immunoblotting. Western blot analysis was also performed using serum samples from COVID-19 cases, Dengue cases, MI cases, healthy individuals and commercial human serum (CHS) (Sigma- 118K0494). Whole-cell lysates and serum sample dilutions (1:10) were prepared and proteins were separated by 10% SDS-PAGE, run at 80 V until optimal resolution was achieved. Proteins were subsequently transferred onto a nitrocellulose membrane using a semi-dry transfer system (20 V, 18 minutes). Following transfer, membranes were blocked in 5% (w/v) skimmed milk solution (HiMedia) prepared in 1× TBST (Tris-buffered saline with 0.1% Tween-20) for 1 hour at room temperature to prevent non-specific binding. After blocking, membranes were washed and incubated overnight at 4°C with primary antibodies diluted in blocking buffer. The following primary antibodies were used: anti-Spike S1 (Abcam, ab275759), anti-GRP78 (Abcam, ab21685), anti-GAPDH (CST, 8884), anti-Beta-Tubulin (CST, 2146) at optimized dilutions.The next day, membranes were washed three times with 1× TBST and incubated with appropriate HRP-conjugated secondary Ab (Goat anti rabbit mAB,, ab97051) for 1 hour at room temperature. After incubation, the membranes were washed again (3× with 1× TBST), and immunoreactive bands were visualized using the Bio-Rad imaging system.

#### Quantitative Real-Time PCR (qRT-PCR)

Total RNA was extracted from 200 µL of cell lysate per well using a standard RNA isolation protocol. The concentration and purity of the RNA were assessed using a NanoDrop One spectrophotometer (Thermo Fisher Scientific). For gene expression analysis, SYBR Green-based one-step quantitative reverse transcription PCR (qRT-PCR) was performed using the Luna® Universal One-Step RT-qPCR Kit (New England Biolabs), following the manufacturer’s instructions. Equal amounts of RNA from each experimental condition were used to ensure consistency across samples.

qRT-PCR reactions were carried out on a QuantStudio™ 5 Real-Time PCR System (Applied Biosystems, Thermo Fisher Scientific). The relative expression levels of target genes were normalized to the housekeeping gene RPL7. Primer sequences used in this study are listed below:

[utbl1]

## 3. RESULTS

### 3.1. Comparative Analysis of Circulating Immune Complexes (CIC) in COVID-19 and Dengue Infections

We aimed to determine if there is any variation in the levels of CICs binding to functional C1q (CIC-C1q) among affected and healthy individuals. We observed a distinct increase in CIC-C1q levels in COVID-19 cases when compared with healthy controls (Fig. 1a). Of the 52 COVID-19 samples analyzed, 13 samples (25%) from both critical and non-critical cases, tested positive in CIC-C1q ELISA. Notably, the prevalence of CIC-C1q was higher in severe COVID-19 cases, with 31.6% (6/19) of these samples (severe COVID-19 cases) testing positive. Our findings also revealed that similar to COVID-19 cases, Dengue cases exhibited a prominent increase in CIC-C1q levels. Of the 19 serum samples studied, 8 samples (42.1%) tested positive for CIC-C1q. In contrast, the apparently healthy controls showed minimal presence of CICs, with only one of 11 apparently healthy samples (9.09%) testing positive (Table 1a).

**Figure 1.**
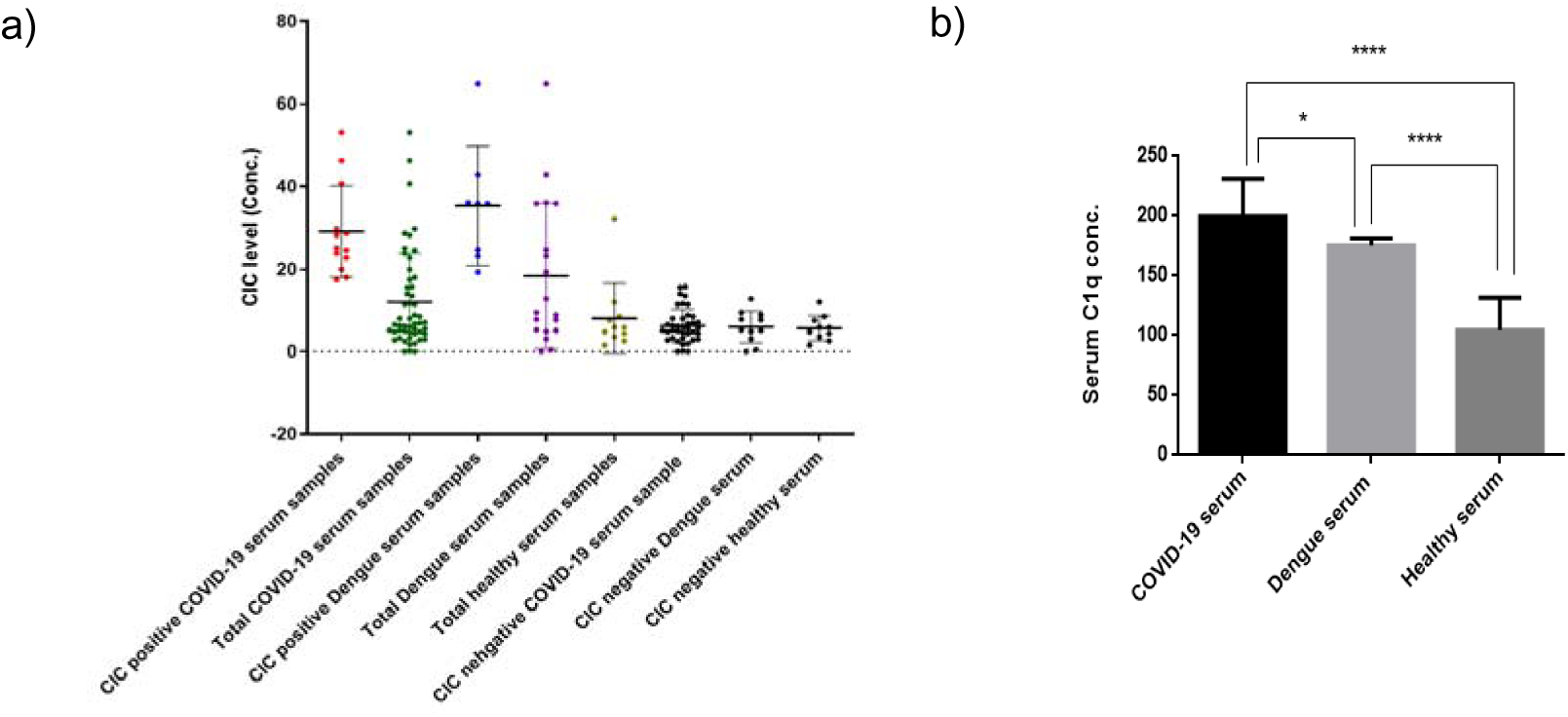
Distribution of CIC-C1q and average concentration of C1q in serum samples from individuals with COVID-19 or Dengue and healthy controls. **(a)** Distribution of CIC-C1q concentrations across the three groups. **(b)** The average C1q concentrations were slightly higher in COVID-19 (n=18) (199.7 ± 31.2 µg/m, SEM = 7.4) serum samples than Dengue (n=13) (p<0.05, *), while healthy individuals (n=10) showed significantly lower levels (P<0.0001, ****) (unpaired Student’s t test) suggesting elevated C1q levels in response to these viral infections. Each data point represents an individual serum sample (biological replicate) for (a); ELISA assays were performed in duplicate (technical replicates) for (a) and (b), and mean values were used for analysis.

**Table 1a.** CIC-Cq positive serum samples.

| Sample type | Sl no. | Manuscript name | Sex | Severity | Concentration | Remarks | CIC conc. (mean) |
| --- | --- | --- | --- | --- | --- | --- | --- |
| COVID-19 serum samples | 1 | C1 | F | STD | 23.758 | (+) | 29.05 |
|  | 2 | C4 | M | STD | 46.264 | (+) |  |
|  | 3 | C9 | M | STD | 28.151 | (+) |  |
|  | 4 | C10 | F | STD | 28.6 | (+) |  |
|  | 5 | C12 | M | STD | 53.102 | (+) |  |
|  | 6 | C14 | M | STD | 19.925 | (+) |  |
|  | 7 | C28 | M | ICU | 17.367 | (+) |  |
|  | 8 | C29 | M | HDU | 29.692 | (+) |  |
|  | 9 | C35 | F | ICU | 17.959 | (+) |  |
|  | 10 | C37 | M | HDU | 22.696 | (+) |  |
|  | 11 | C41 | M | STD | 24.529 | (+) |  |
|  | 12 | C42 | M | ICU | 25.009 | (+) |  |
|  | 13 | C46 | M | HDU | 40.652 | (+) |  |
| Dengue serum samples | 1 | D6 | F | STD | 19.249 | (+) | 35.33 |
|  | 2 | D9 | M | STD | 35.856 | (+) |  |
|  | 3 | D11 | M | STD | 36.067 | (+) |  |
|  | 4 | D12 | F | STD | 42.91 | (+) |  |
|  | 5 | D13 | M | STD | 64.973 | (+) |  |
|  | 6 | D14 | F | STD | 35.865 | (+) |  |
|  | 7 | D15 | M | STD | 23.125 | (+) |  |
|  | 8 | D16 | F | STD | 24.601 | (+) |  |
| Healthy serum sample | 1 | H3 | M | STD | 32.154 | (+) | 32.154 |

Table 1a shows CIC-C1q positive serum samples (with a cut-off value of 16 units), while table 1b displays CIC-C1q negative serums across COVID-19, Dengue, and healthy samples. Of the 52 COVID-19 cases analyzed (30 males and 22 females), 19 individuals (36.5%) presented with severe disease requiring ICU or HDU care. CIC-C1q positivity was observed in 13 individuals (25%), comprising 10 males (19.2%) and 3 females (5.8%). Of these 13 positive samples, 6 (46.2%) belonged to the severe category (ICU/HDU). In all severe cases of COVID-19, 32% (6 out of 19) serum samples are CIC-C1q positive. The mean CIC-C1q concentration in positive COVID-19 samples was 29.1 ± 11.0 (SD), with a standard error (SE) of 3.05, while the negative group showed a mean of 6.2 ± 4.1 (SD), SEM = 0.7.

**Table 1b.** CIC-C1q negative serum samples. Table 1a, b. Circulating Immune Complex (CIC-C1q) levels and distribution in serum samples from COVID-19, Dengue, and healthy individuals.

| Sample type | Sl no. | Manuscript name | Sex | Severity | Concentration | Remarks | CIC conc. mean |
| --- | --- | --- | --- | --- | --- | --- | --- |
| COVID-19 serum samples | 1 | C2 | F | STD | 8.015 | (-) | 6.20 |
|  | 2 | C3 | M | STD | 5.498 | (-) |  |
|  | 3 | C5 | F | STD | 6.7 | (-) |  |
|  | 4 | C6 | F | STD | 6.287 | (-) |  |
|  | 5 | C7 | F | STD | 7.977 | (-) |  |
|  | 6 | C8 | M | STD | 3.206 | (-) |  |
|  | 7 | C11 | M | STD | 2.454 | (-) |  |
|  | 8 | C13 | M | STD | 11.547 | (-) |  |
|  | 9 | C15 | F | STD | 6.888 | (-) |  |
|  | 10 | C16 | F | STD | 4.333 | (-) |  |
|  | 11 | C17 | F | STD | 7 | (-) |  |
|  | 12 | C18 | F | STD | 13.463 | (-) |  |
|  | 13 | C19 | F | ICU | 6.522 | (-) |  |
|  | 14 | C20 | M | HDU | 2.955 | (-) |  |
|  | 15 | C21 | M | ICU | 5.159 | (-) |  |
|  | 16 | C22 | M | STD | 2.723 | (-) |  |
|  | 17 | C23 | F | STD | 5.101 | (-) |  |
|  | 18 | C24 | M | HDU | 11.366 | (-) |  |
|  | 19 | C25 | F | STD | 8.813 | (-) |  |
|  | 20 | C26 | M | HDU | 5.565 | (-) |  |
|  | 21 | C27 | F | STD | 6.145 | (-) |  |
|  | 22 | C30 | F | STD | 0 | (-) |  |
|  | 23 | C31 | M | STD | 13.995 | (-) |  |
|  | 24 | C32 | M | STD | 3.216 | (-) |  |
|  | 25 | C33 | M | STD | 4.763 | (-) |  |
|  | 26 | C34 | M | STD | 1.923 | (-) |  |
|  | 27 | C36 | M | HDU | 4.612 | (-) |  |
|  | 28 | C38 | F | HDU | 4.974 | (-) |  |
|  | 29 | C39 | M | ICU | 4.306 | (-) |  |
|  | 30 | C40 | M | STD | 8.514 | (-) |  |
|  | 31 | C43 | F | ICU | 0.069 | (-) |  |
|  | 32 | C44 | M | STD | 1.726 | (-) |  |
|  | 33 | C45 | M | HDU | 2.773 | (-) |  |
|  | 34 | C47 | F | ICU | 11.355 | (-) |  |
|  | 35 | C48 | F | HDU | 15.575 | (-) |  |
|  | 36 | C49 | M | STD | 6.005 | (-) |  |
|  | 37 | C50 | F | HDU | 0 | (-) |  |
|  | 38 | C51 | F | STD | 15.713 | (-) |  |
|  | 39 | C52 | M | STD | 4.758 | (-) |  |
| Dengue serum samples | 1 | D1 | M | STD | 7.714 | (-) | 5.93 |
|  | 2 | D2 | M | STD | 8.879 | (-) |  |
|  | 3 | D3 | F | STD | 5.122 | (-) |  |
|  | 4 | D4 | M | STD | 9.48 | (-) |  |
|  | 5 | D5 | F | STD | 0.388 | (-) |  |
|  | 6 | D7 | F | STD | 7.79 | (-) |  |
|  | 7 | D8 | M | STD | 3.055 | (-) |  |
|  | 8 | D10 | M | STD | 4.859 | (-) |  |
|  | 9 | D17 | M | STD | 5.238 | (-) |  |
|  | 10 | D18 | F | STD | 12.705 | (-) |  |
|  | 11 | D19 | F | STD | 0 | (-) |  |
| Healthy serum samples | 1 | H1 | F | STD | 3.51 | (-) | 5.63 |
|  | 2 | H2 | F | STD | 5.885 | (-) |  |
|  | 3 | H4 | M | STD | 5.814 | (-) |  |
|  | 4 | H5 | F | STD | 4.36 | (-) |  |
|  | 5 | H6 | M | STD | 8.473 | (-) |  |
|  | 6 | H7 | M | STD | 4.75 | (-) |  |
|  | 7 | H8 | F | STD | 2.552 | (-) |  |
|  | 8 | H9 | M | STD | 11.937 | (-) |  |
|  | 9 | H10 | F | STD | 1.515 | (-) |  |
|  | 10 | H11 | F | STD | 7.564 | (-) |  |

In the Dengue group (n = 19), 8 samples (42.1%) were CIC positive, with a mean concentration of 35.3 ± 14.4 (SD), SEM = 5.1, compared to 5.9 ± 3.9 (SD), SEM = 1.2 in the CIC-C1q negative Dengue serum samples.

Among healthy individuals (n = 11), only one sample (9.1%) was CIC-C1q positive, while the rest are CIC negative with a mean of 5.6 ± 3.1 (SD), SEM = 1.0.

Each data represents an individual serum sample (biological replicate); ELISA assays were performed in duplicate (technical replicates), and mean values were used for analysis.

CIC-C1q=Circulating Immune Complex; ICU=Intensive Care Unit; HDU=High Dependency Unit; SD=Standard Deviation; SE=Standard Error of the mean

### 3.2 Activation of C1q-mediated Complement System in Individuals with severe COVID-19 and Dengue Infection

Excessive C1q deposition has been reported to be linked to thrombotic issues, atherosclerotic advancement, and cardiovascular complications^38,39^. In order to determine the degree of C1q-facilitated activation of the classical complement pathway and thrombotic problems, C1q ELISA was implemented to quantify the plasma levels of soluble C1q proteins in a set of healthy as well as affected serum samples. All the CIC-C1q-positive serum samples and a sample set of CIC-C1q negative serum samples were analyzed across the affected and control groups. The mean C1q levels showed a marked increase in both COVID-19 and Dengue cases when compared to healthy controls.

Contrary to healthy samples, a significant increase of mean C1q levels at 199.7 μg/ml (P<0.0001) for COVID-19 and 175.5 μg/ml (P<0.0001) for the Dengue serum samples, were observed (Table 2; Fig. 1). These levels were higher than the mean C1q level of the apparently healthy serums (104.7) (this study) and even the reference level for C1q (113±40 μg/ml) in healthy serums^40^.

**Table 2.**
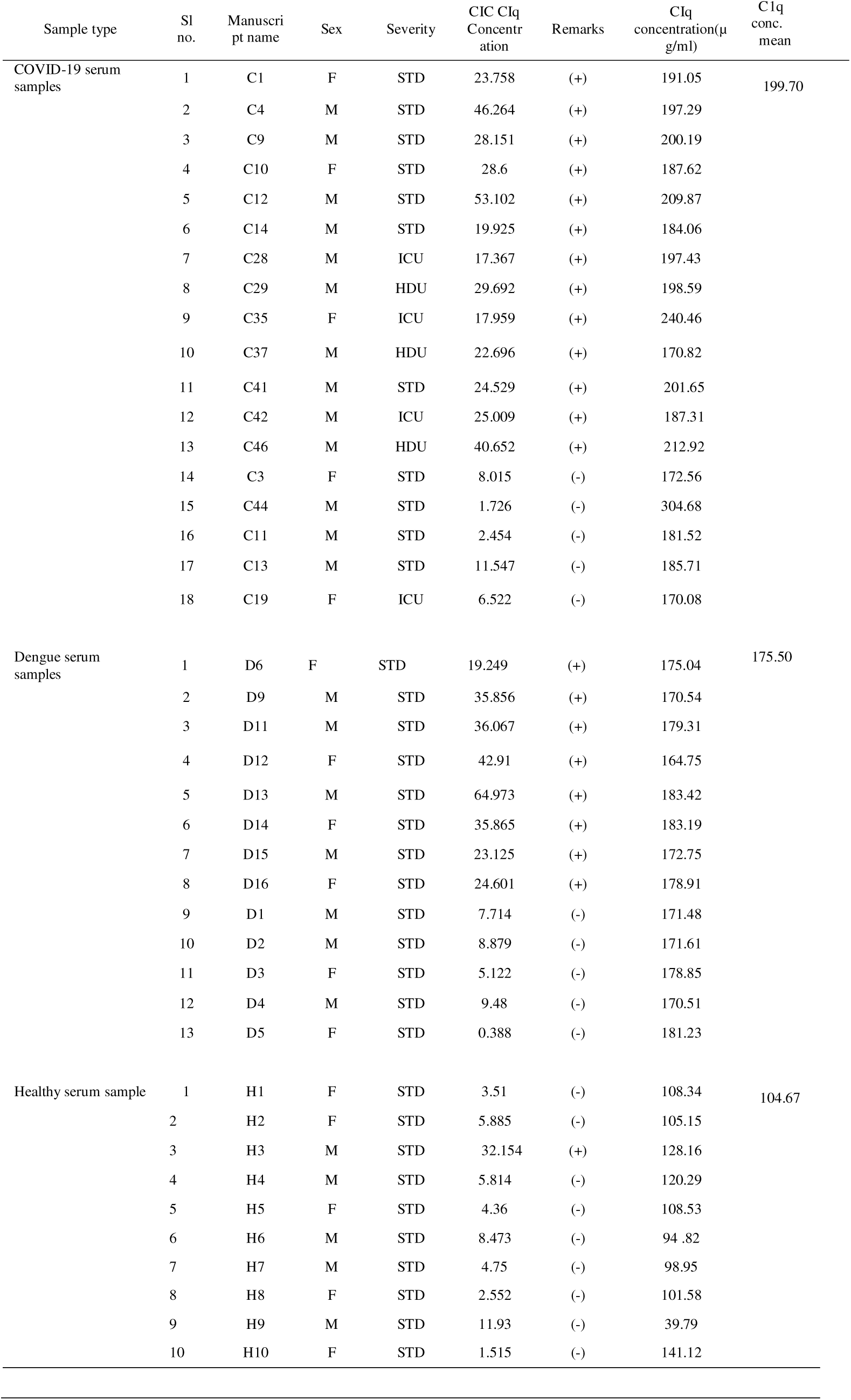
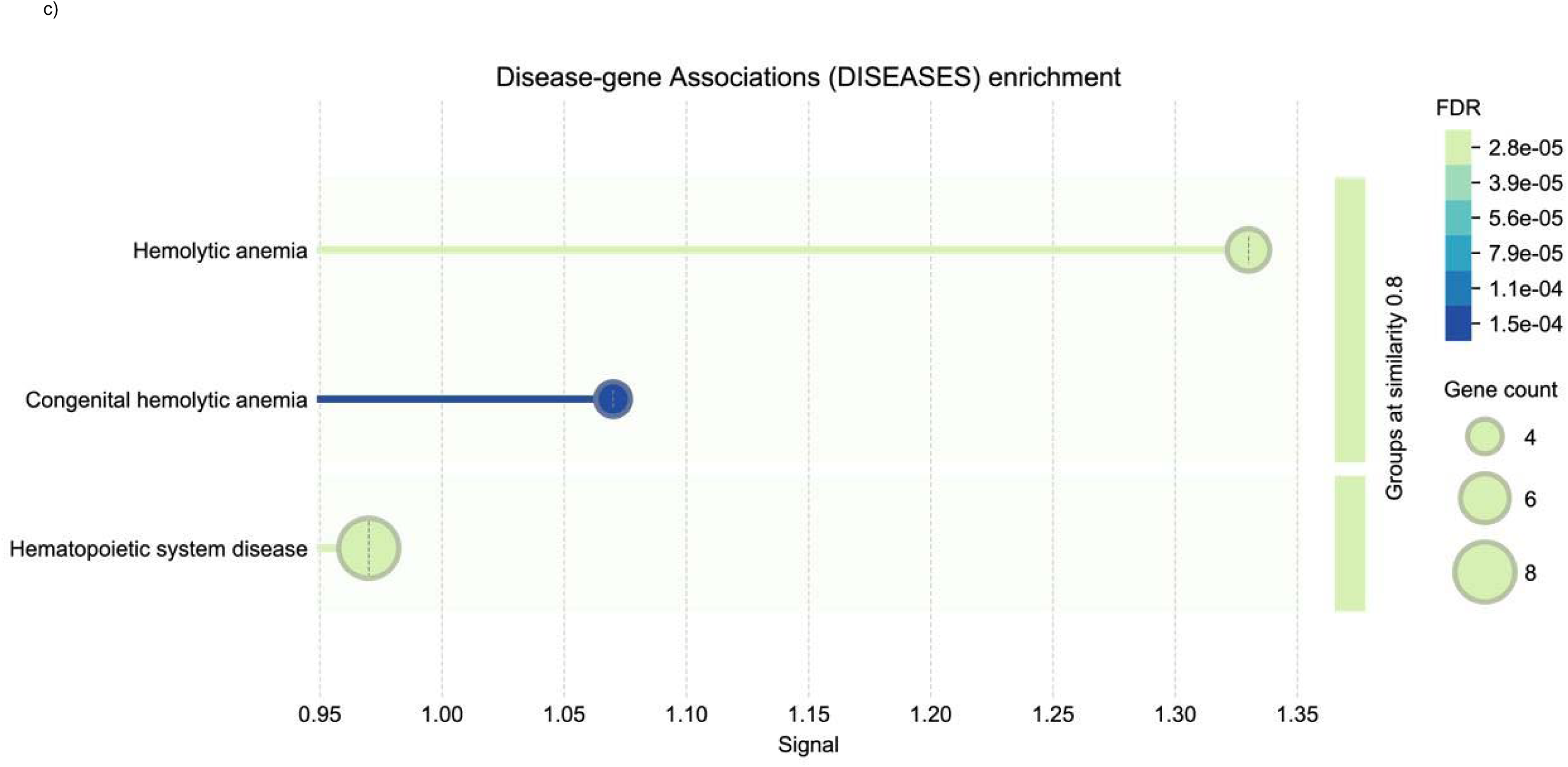
Serum C1q levels in healthy controls, COVID-19 and Dengue serum samples, stratified by CIC-C1q status. C1q concentrations were measured by C1q ELISA in the serum samples. Mean C1q levels were significantly elevated in COVID-19 (n=18) (199.7 ± 31.2 µg/ml, SEM = 7.4) and Dengue cases (n=13) (175.5 ± 5.7 µg/ml, SEM = 1.6) compared to healthy controls (n=10) (104.7 ± 26.88 µg/ml, SEM = 8.49), with p < 0.0001, **** (unpaired Student’s t-test). Subgroup analysis based on CIC-C1q status showed no significant difference in mean C1q levels between CIC-C1q positive and negative samples within each disease group. One healthy control serum sample was CIC-C1q positive and had a C1q concentration of 128.2 µg/ml. Each data point represents an individual serum sample (biological replicate); ELISA assays were performed in duplicate (technical replicates), and mean values were used for analysis.

These findings suggest activation of the classical complement pathway in both COVID-19 and Dengue infections. Given the role of C1q complement protein in immune complex–mediated complement activation, this elevation may reflect enhanced inflammatory responses and immune complex formation in infected individuals.

### 3.3. Proteomics analysis of serum samples from COVID-19 and Dengue Infections

Given the observed elevation of C1q complement protein and circulating immune complexes (CIC)–C1q levels in infected samples, indicative of activation of the classical complement pathway, it was interesting to assess the serum proteomic profile of these two significant viral infections to obtain a global view on enrichment of possible risk factors (due to the infections) that could lead to cardiovascular complications.

To investigate the key proteomic changes in individuals suffering from either Dengue or COVID-19, an ESI/nano-LC-MS/MS analysis was carried out on an LTQ Orbitrap XL, on aset of serum samples from the affected and healthy individuals (Supplementary table 3). The mass spectra acquired from the 145 minutes LC-MS/MS run-on analysis using Proteome Discoverer v1.4.0 revealed the identification of 171, 178, and 175 annotated proteins from depleted plasma samples of Groups I (COVID-19 cases), II (Dengue cases), and III (healthy controls), respectively. Functional classification of the proteome of serum samples from COVID-19 and Dengue individuals broadly revealed enrichment of immune response pathways.

#### Comparative analysis of protein expression of COVID-19, Dengue and healthy serums based on different categories

Healthy individuals with no prior or current history of COVID-19 or Dengue infection were used as controls and served as the reference group for comparative serum proteomic analyses. Differentially expressed proteins identified in the COVID-19 (Supplementary Fig. 1a–d) and Dengue (Supplementary Fig. 2a–d) cohorts were annotated and functionally classified using Gene Ontology (GO) categories – including biological processes, molecular functions, protein classes, and pathways through the PANTHER database (v19.0) (Mi et al., 2021).

Proteins commonly identified in both infection groups spanned diverse functional classes, including defense and immunity proteins, transport and carrier proteins, extracellular matrix components, and structural proteins. Pathway analysis revealed enrichment of multiple immune- and inflammation-associated pathways, including blood coagulation, B cell activation, T cell activation, Toll-like receptor signaling, chemokine- and cytokine-mediated inflammation, VEGF signaling, and AT-II signaling via G proteins and β-arrestins. Notably, pathways related to immune activation and coagulopathy, such as B cell activation (P00010),

T cell activation (P00053), Toll-like receptor signaling (P00054), and blood coagulation (P00011), were significantly over-represented in both infection groups.

Despite distinct pathways and functions associated with COVID-19 and Dengue, both conditions exhibit overlapping clinical manifestations, which may be part attributable to shared molecular responses. To evaluate the extent of proteomic overlap, protein lists from both infection groups were compared using InteractiVenn^41^. This analysis revealed that 59.9% of identified proteins were common to both infections, whereas 18.4% were uniquely detected in the COVID-19 serum proteome (Fig. 2a).

**Figure 2.**
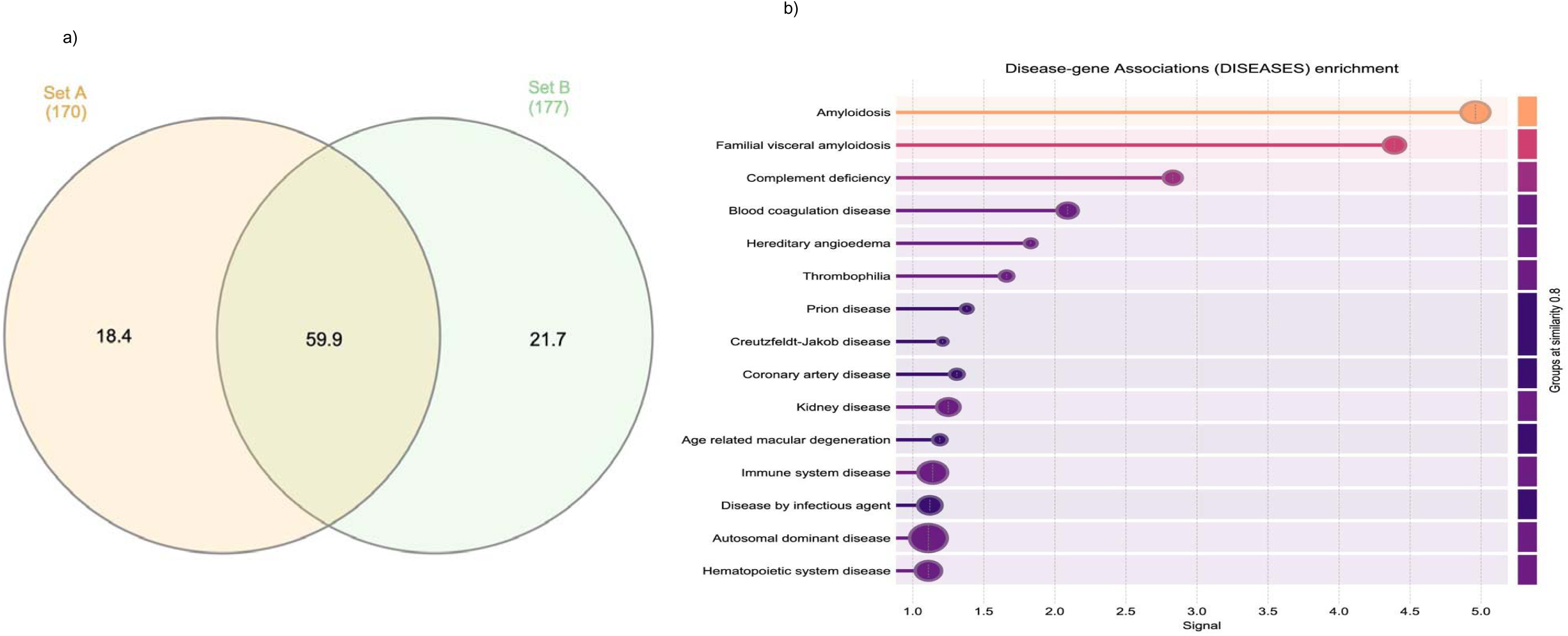
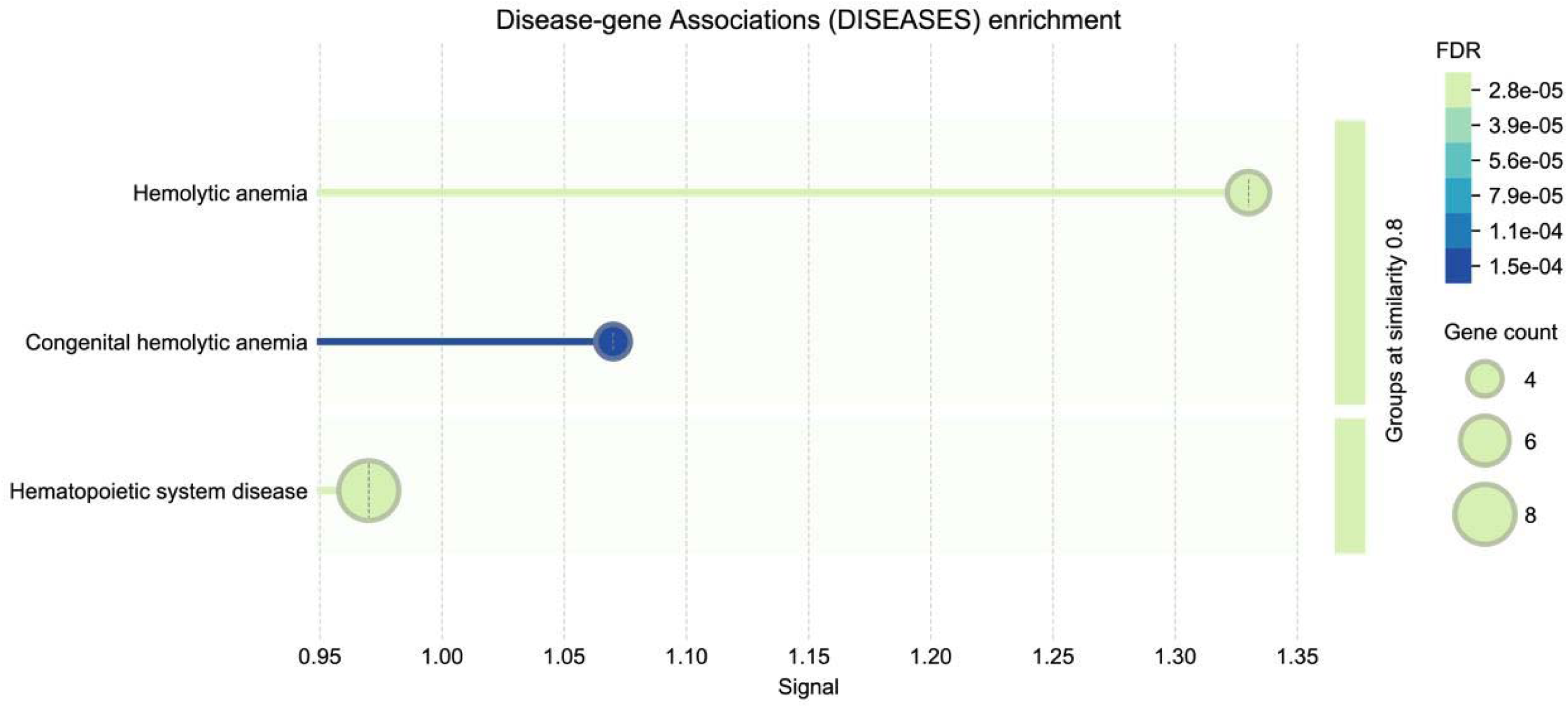
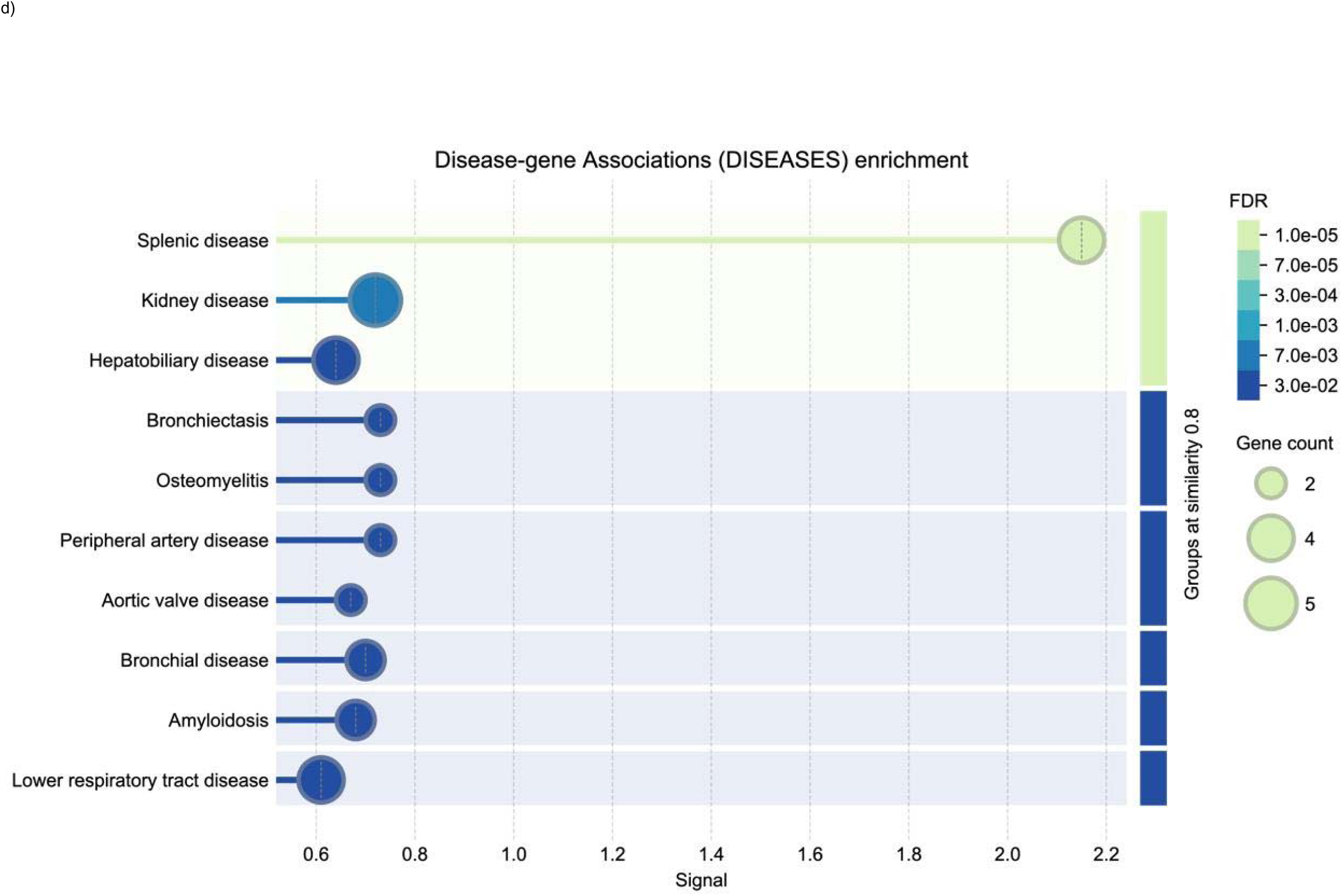
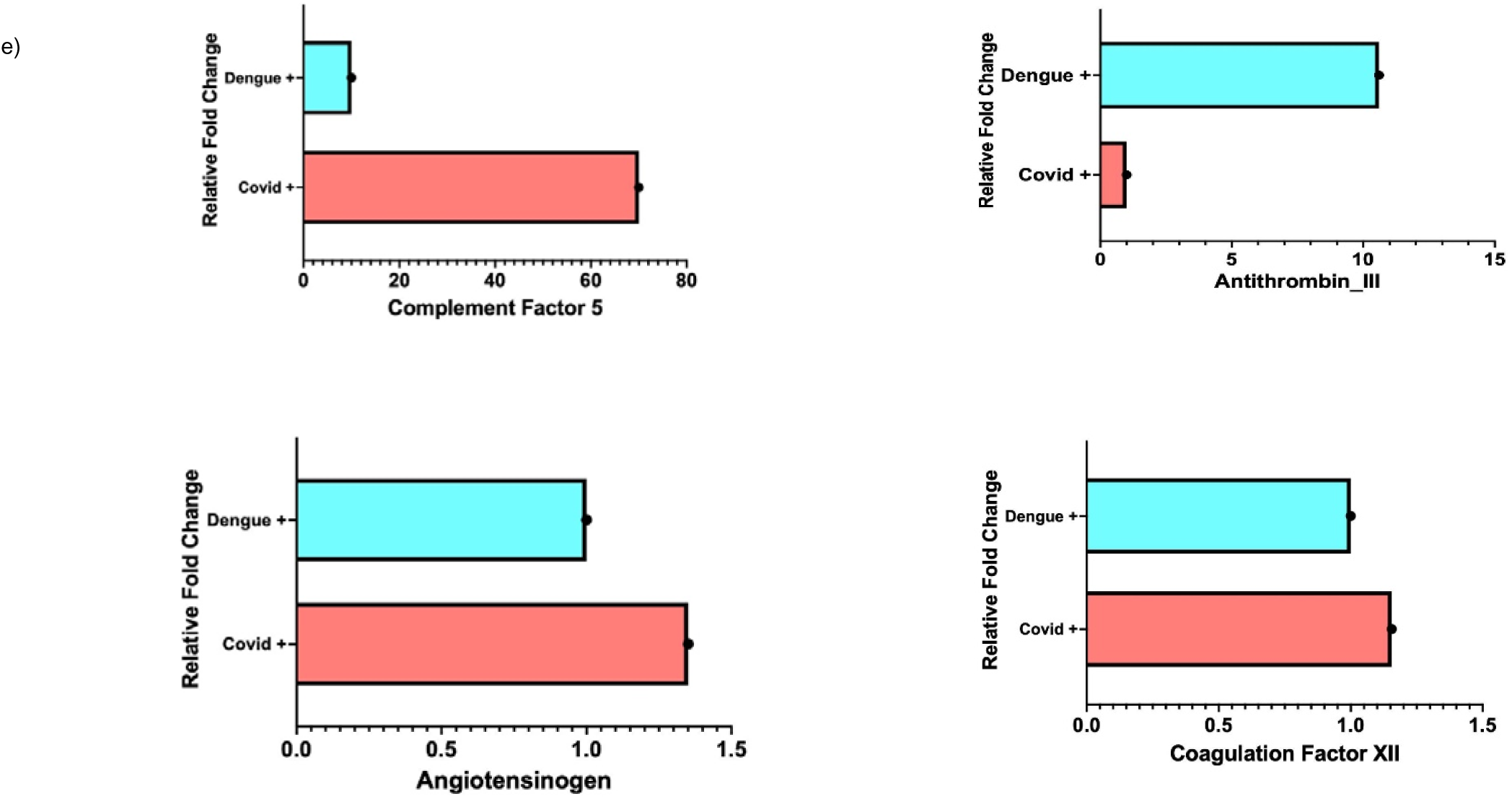
Comparative Proteomic Analysis of Serum from COVID-19 and Dengue cases. (a) According to the web-based analytic tool ractiVenn, 59.9% of the proteins were shared by the two groups. However, 18.4% and 21.7% of the proteins were specific to COVID-19 and ngue, respectively. (b) There were similarities between the identified proteins from the mass spectrometry data of serum samples from both ction groups. These proteins were linked to a number of different disorders. These include immune system problems, kidney and diovascular issues and conditions affecting coagulation factors and the complement cascade. (c) From Disease-gene association enrichment lysis, it was found that Dengue cases presented the risk of developing hemolytic anemia or diseases of the hematopoietic system and (d) VID-19 cases were more susceptible to thrombus formation, which increased their risk of developing diseases such as aortic valve disease or ipheral artery disease. (e) Angiotensinogen, complement factor 5, and coagulation factor XII were upregulated in COVID-19, whereas ithrombin was downregulated in COVID-19 compared to Dengue samples.

#### Disease-gene association studies of the proteins revealed that cardiovascular risks were higher in COVID-19 cases

As stated earlier, both COVID-19 and Dengue cases were at a significant risk of developing health complications. Therefore, it was crucial to analyze the specific types of health issues they are more likely to face. To have a comprehensive overview of how the annotated proteins in our list can be linked to incidences of metabolic or lifestyle diseases at a molecular level, we conducted gene-disease association studies using the DISEASE software.

The predicted diseases were sorted based on the signal attribute (minimum signal value set at ≥ 0.01); defined as a weighted harmonic mean between the observed/expected ratio and -log [False Discovery Rate (FDR)]. The maximum FDR score was fixed at ≤ 0.05. As observed, the annotated proteins from the mass spectrometry data of serum samples from both infection groups showed commonalities in proteins associated with various other conditions. These included diseases of the immune system, kidney and cardiovascular complications, as well as disorders involving the complement cascade and coagulation factors, to name a few (Fig. 2b). Although individuals from both infection groups were at an increased risk of developing cardiovascular issues, the types of cardiac complications differed between the two. Dengue cases were more prone to bleeding, which elevated their risk of developing hemolytic anemia or diseases of the hematopoietic system (Fig. 2c). In contrast, COVID-19 cases were more likely to develop thrombi, putting them at greater risk for conditions like peripheral artery or aortic valve diseases (Fig. 2d).

To further predict and identify all possible protein-protein interactions (PPIs) unique to COVID-19 infection associated proteome, we used a web resource tool and biological database (STRING) that is considered a gold benchmark for investigating the interactions between genes that are commonly dysregulated. As indicated from the enrichment analysis (PPI enrichment p-value: 1.11e-16) the proteins uniquely dysregulated in COVID-19 infection were at least partially connected, as a biological group. About 37 different protein sub-clusters/protein associations among 19 proteins were predicted using the STRING database (Supplementary Fig. 3). These biological groups can be further sub-classified into two major local networking clusters namely the high-density lipoproteins; and the complement and coagulation cascades and protein-lipid complex (Supplementary Fig. 4).

#### COVID-19 serum samples had elevated levels of key proteins linked to coagulopathy and vasoconstriction

Although the networking clusters were somewhat similar in both infection groups, we wanted to determine if there were statistically significant differences in abundance of proteins between COVID-19 and Dengue population samples. To conduct the standard two-population differential analyses, we used SIEVE^TM^ software to compare the two sample files. As expected from ratio analysis, proteins that are involved in coagulation system activation and platelet aggregation such as coagulation factor XII, complement factor 5 (C5) and Angiotensinogen were up-regulated in COVID-19.

The factor XII-driven system initiates inflammatory and coagulation processes through the bradykinin-producing kallikrein-kinin system and the intrinsic coagulation pathway, respectively^42^. Inflammation and thrombosis are linked by thrombin activation of C5. Increased C5 plasma levels serve as a novel circulating biomarker of subclinical atherosclerosis^43,44^. Finally, angiotensinogen activation and the subsequent synthesis of AT-II are believed to be a cause of cardiotropin-1-induced cardiac myocyte hypertrophy. Also, AT-II type-1 receptor (AT1R)’s positive feedback control of angiotensinogen expression could contribute to further vasoconstriction^45,46,47^. Antithrombin-III was observed to be down-regulated in COVID-19, compared to Dengue samples (Fig. 2e). Antithrombin-III (AT-III) is a plasma protein that inhibits several activated pro-coagulants^48^.

### 3.4. Impact of SARS-CoV-2 spike gene transfection in different cell lines including cardiomyocytes upon spike expression

Besides the lungs, the SARS-CoV-2 virus has been shown to harm other organs. Remarkably, the blood of COVID-19 has been reported to contain circulating SARS-CoV-2 spike (S) protein, including in some studies, following infection and mRNA vaccination^47^. It has been found that circulating S protein can bind to receptors, potentially contributing inflammation and damage to cells, tissues, and organs^50^. However, the exact relationship between spike protein, inflammation, tissue damage, and coagulopathy are still to be further elucidated. Therefore, different cell lines including cardiomyocytes were transfected with spike gene and the cells were subsequently analyzed for cellular pathology.

#### 3.4.1. The SARS-CoV-2 spike antibody showed cross-reactivity with a different protein (not spike protein) in heart and lung cell lines, as per immunoblotting analysis

Western blot was carried out on heart (H9c2), lung (A549), and kidney (HEK293T) cell lysates, using SARS-CoV-2 S1 spike Ab. As shown in the blot (Fig. 3; Supplementary figure 7), HEK293T (human embryonic kidney-derived) cells transfected with spike protein, displayed a clear band between 100–130 kDa, consistent with the molecular weight of the S1 spike protein (∼120 kDa). This band was absent in the corresponding un-transfected control lysates. In contrast, lysates from A549 (human alveolar basal epithelial cell line derived) and H9c2 (embryonic rat heart tissue derived) cells did not show the expected spike band; instead, a prominent band was observed between 70-100 kDa, suggesting Ab cross-reactivity with an endogenous cellular protein. These results indicated that the S1 Ab can indeed, bind to a non-spike protein in heart and lung-derived cell lines.

**Figure 3:**
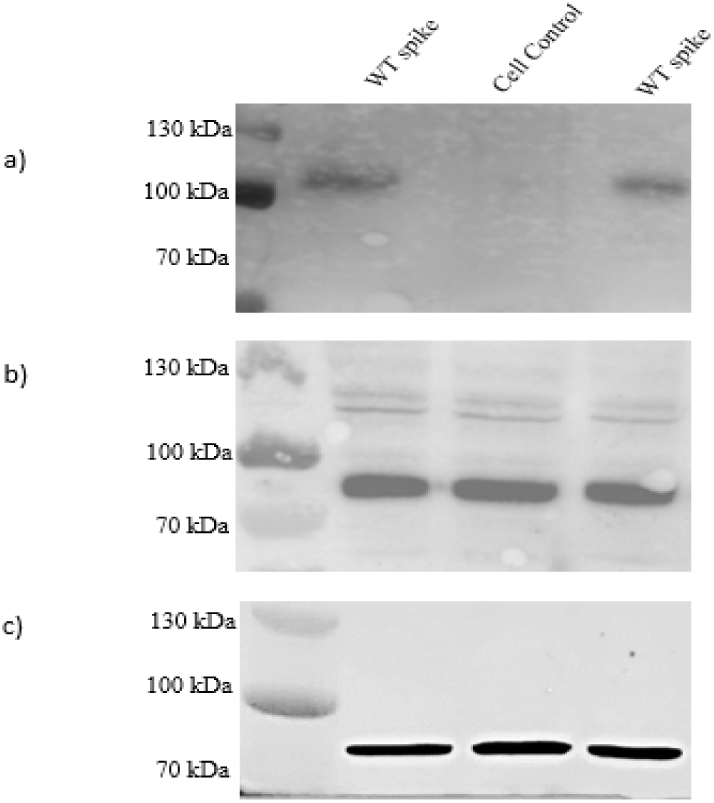
Analysis by immunoblotting showed that the spike antibody used for SARS-CoV-2 detection cross-reacts with a different protein in cardiac and pulmonary cell lines.

Western blotting was performed on lysates from spike-transfected cells collected 72 hours post-transfection. Panel (a) shows HEK293T (kidney) cell lysate, (b) shows H9c2 (heart) cell lysate, and (c) shows A549 (lung) cell lysate. A distinct band between 100–130 kDa, corresponding to the S1, was visible only in case of HEK293T cells (a). In H9c2 and A549 lysates (b, c), no spike band was detected; instead, a non-specific band near 70 kDa was observed, indicating possible cross-reactivity of the S1 Ab with an endogenous protein in these cell lines. All experiments were performed with three independent biological replicates.

WT= Wild Type

#### 3.4.2. GRP78 was identified as the protein cross-reacting with the SARS-CoV-2 S1 spike Ab in heart and lung cell lines

In order to identify the protein corresponding to the 70-100 kDa band observed in A549 and H9c2 cell lysates during immunoblotting, the excised gel region (70–100 kDa) was subjected to in-gel digestion followed by mass spectrometry analysis. Mascot search results revealed the presence of two candidate proteins within this molecular weight range, namely glucose-regulated protein 78 (GRP78/BiP) and moesin (Fig. 4). GRP78 has a molecular weight of approximately 78 kDa, while moesin is approximately 63.4 kDa. Based on this size difference evident from the observed migration pattern on the blot and the mean Mascot score, GRP78 was identified as the most likely protein recognized by the S1 spike Ab in both A549 and H9c2 cells. These findings suggested that the spike S1 Ab could cross-react with endogenous GRP78 in heart and lung-derived cell lines.

**Figure 4:**
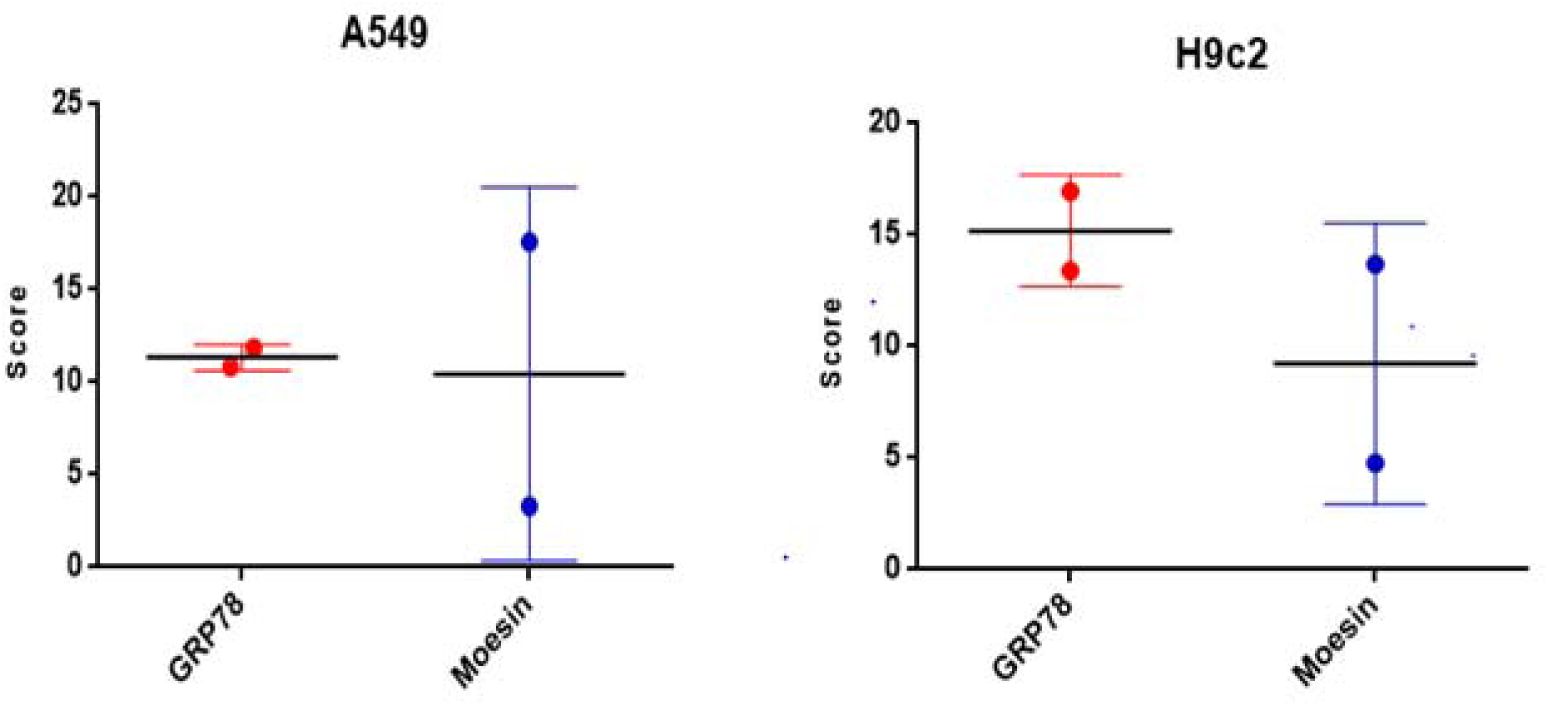
Mass spectrometry analysis of the 70–100 kDa band from A549 and H9c2 cell lysates revealed GRP78 as the likely cross-reactive protein. The excised protein band between 70–100 kDa from SDS-PAGE gels was subjected to in-gel digestion and analyzed by mass spectrometry. Mascot scores of the top-identified proteins were plotted for (a) A549 and (b) H9c2 cell lysates. In both samples, GRP78 (BiP) and moesin were detected with high scores. Given its molecular weight (∼78 kDa) and alignment with the observed band on immune-blotting, GRP78 was found the most likely candidate for the protein bound by the S1 spike Ab in A549 and H9c2 cells.

#### 3.4.3. Distinct spike protein expression patterns observed in three cell lines following transfection

The SARS-CoV-2 pUNO1 spike plasmid was transfected into three different cell lines, and spike protein expression was assessed via immunofluorescence on staining with spike S1 Ab at 48 hours post-transfection. Confocal microscopy revealed three distinct expression patterns across the three cell lines (Fig. 5a), which were further supported by quantitative analysis of mean fluorescence intensity (MFI) (Fig. 5b).

**Figure 5:**
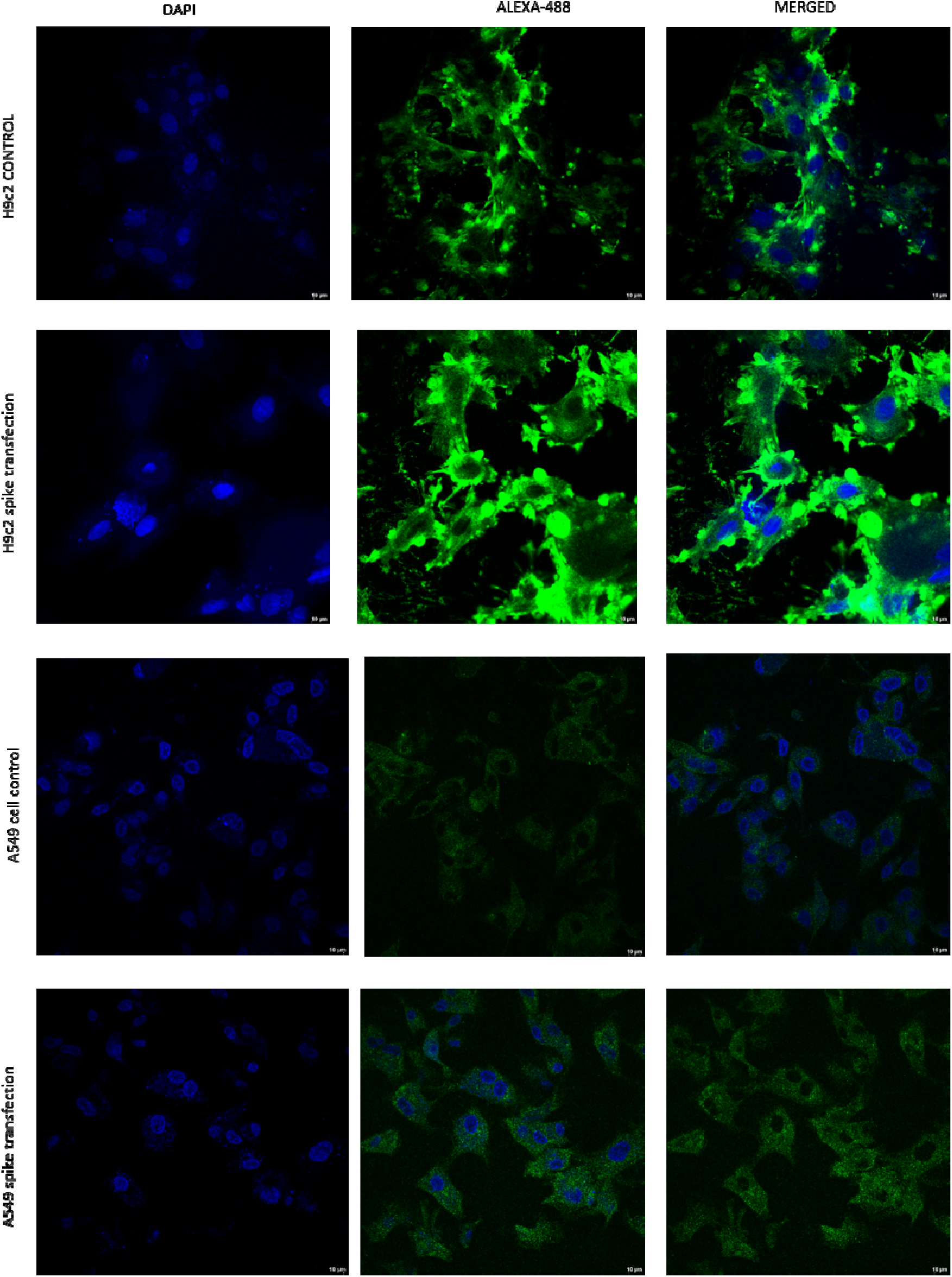
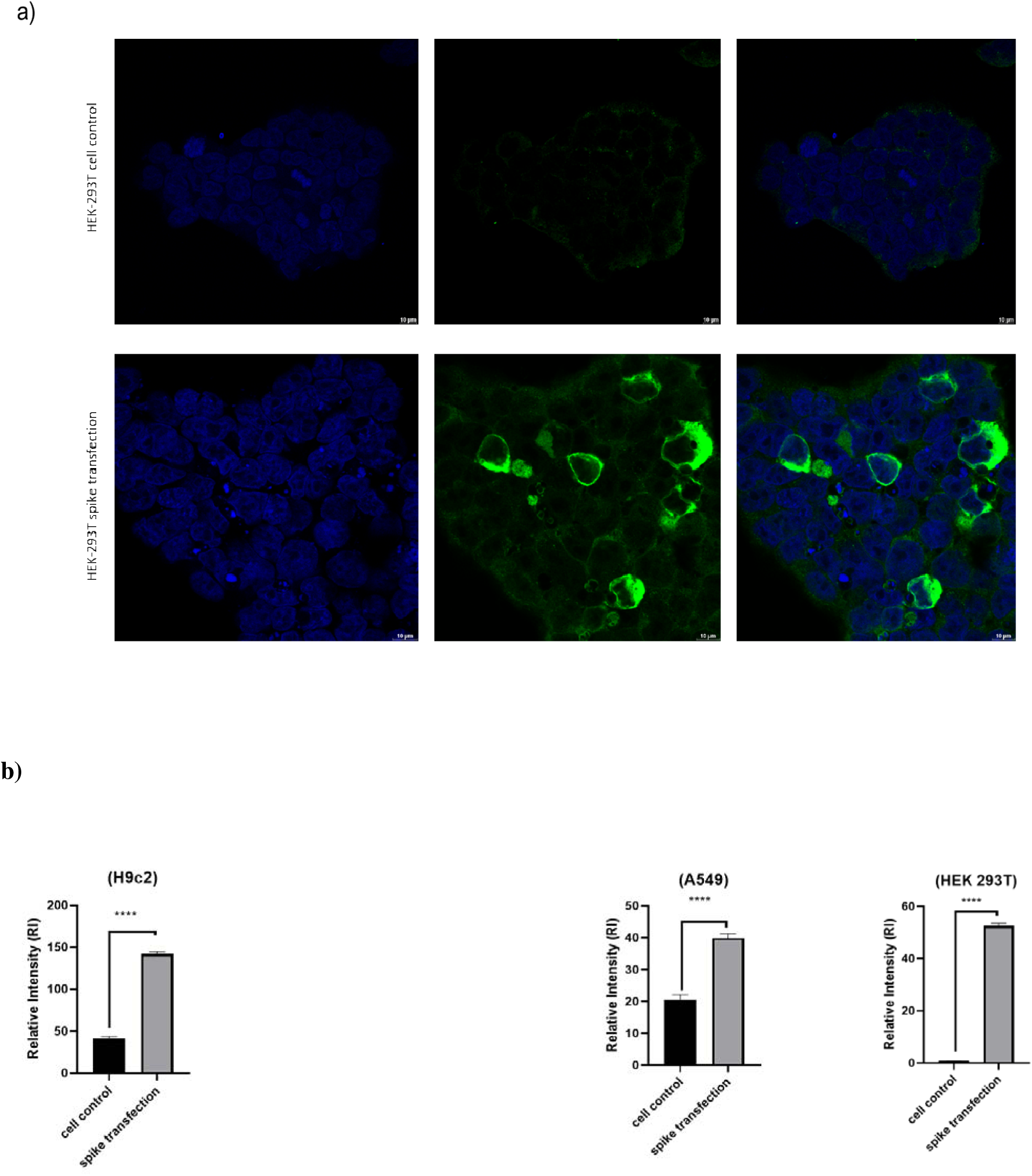
SARS-CoV-2 spike protein expression in different cell lines. H9c2, A549 and HEK293T cells were transfected with pUNO1 spike plasmid and stained with S1 Ab at 48 hours post-transfection. Confocal microscopy revealed distinct expression patterns. (a) H9c2 cells exhibited the highest surface expression with prominent membrane aggregation of spike protein with A549 and HEK293T cells showed moderate and high expression respectively. (b) The graphs represent the mean fluorescence intensity (MFI) of S1 Ab binding per cell (P<0.0001, Data are presented as mean ± SEM (n=3) for all the cell lines).

Among them, H9c2 cells exhibited the highest level of surface expression, with pronounced aggregation of spike protein across the cell membrane. The protein didn’t appear to accumulate prominently in the cytoplasm/endoplasmic reticulum (ER) based on qualitative assessment of confocal images (Fig. 5a) The HEK293 cells showed bright green cytoplasm suggesting accumulation of spike protein in the ER. Notably, the lung-derived cell line displayed a markedly different localization pattern compared to the other two cell types-spike was diffusely spread throughout cytoplasm and did not appear to accumulate within the ER or cell membrane (Fig. 5a).

Background staining of cell controls appeared to be more in case of the lung and cardiac cell lines possibly due to cross-reactivity of S1 Ab with endogenous proteins like GRP78 (Fig. 5b).

#### 3.4.5. Confocal microscopy showed that spike and GRP78 antibodies, both could bind to the same regions in spike gene transfected cardiomyocyte cell line

Using mass spectrometry, it was observed that spike S1 Ab could bind to GRP78 in H9c2 and A549 cells. A co-localization analysis was conducted using confocal microscopy in the H9c2 cell line to further validate this data. It was evident from co-localized staining that the spike S1 Ab and GRP78 Ab both were binding to the same region of the transfected cells (Fig. 6).

**Fig. 6:**
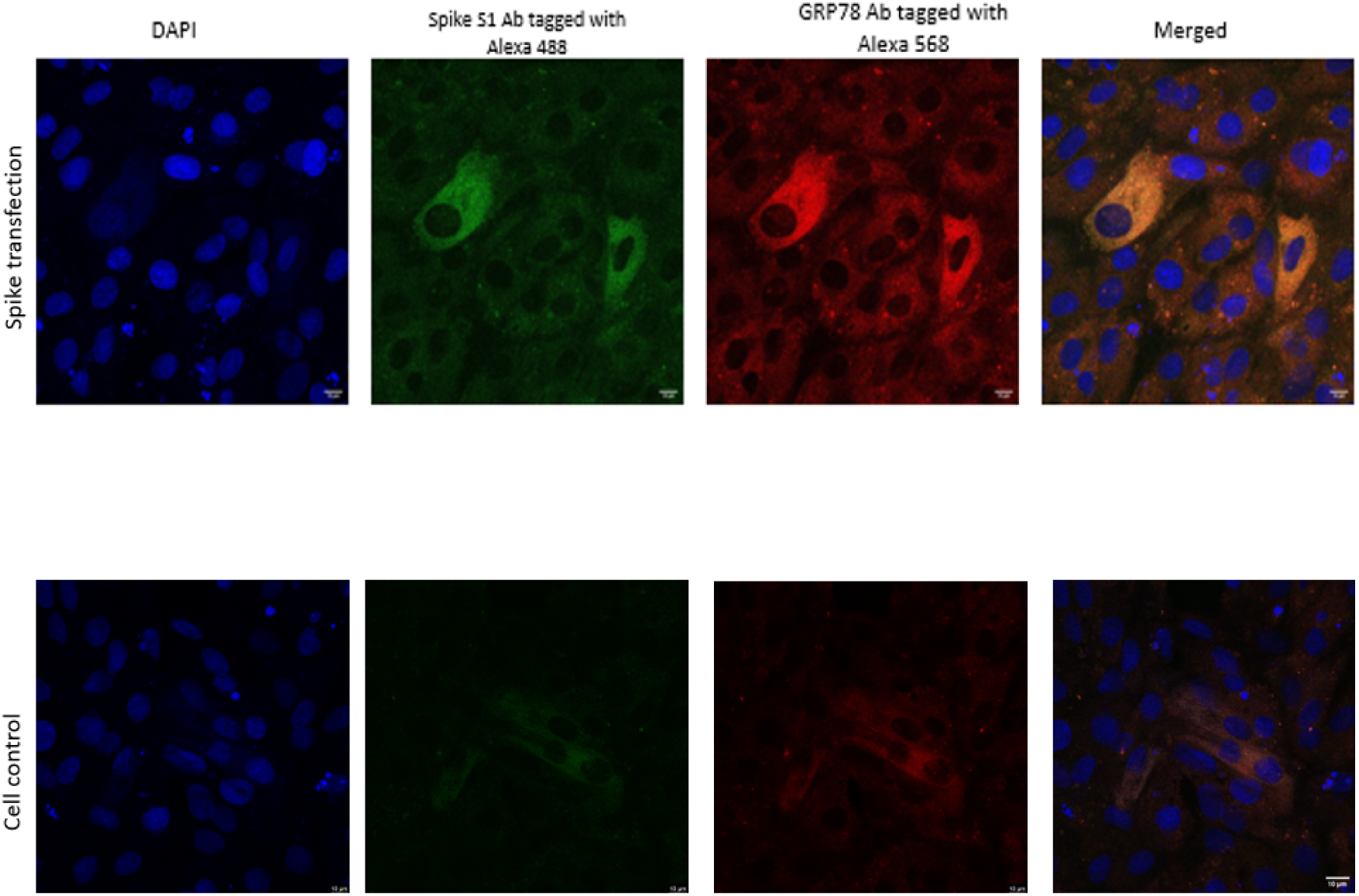
Co-localization of spike protein and GRP78 in spike-transfected cells. Confocal microscopy images of cell control and pUNO1 spike transfected H9c2 cardiomyocyte cells stained with spike S1 Ab (green) and GRP78 Ab (red). Co-localization was observed across the cells, supporting the interaction between spike protein and GRP78.

### 3.5. Increased GRP78 expression at the RNA level, was detected in spike gene-transfected cells

In order to assess the effect of SARS-CoV-2 spike protein expression on GRP78 gene expression, A549 cells were transfected with Puno-1 spike plasmid. The mRNA level of GRP78 was estimated at 48 hours post-transfection. Compared to the un-transfected control group, cells expressing spike protein exhibited nearly two-fold increase in GRP78 mRNA expression (Fig. 7). This observation suggested that expression of the viral spike protein alone was sufficient to induce ER stress in host cells, potentially contributing to the cellular environment associated with COVID-19 pathogenesis. It is plausible that GRP78 gene expression was heightened in response to this stress.

**Figure 7:**
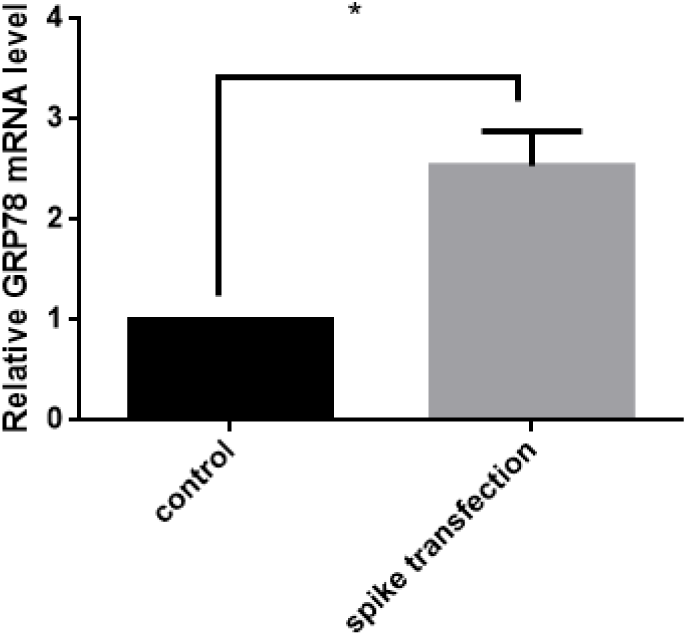
Relative GRP78 mRNA level in cell control and spike plasmid transfected A549 cells. The GRP78 mRNA level in spike gene-transfected cells were almost twice (P<0.05, *) that of the control cells.

#### Increased GRP78 protein expression in spike-expressing cells

Western blot analysis revealed an increase in GRP78 protein level in A549 cells and H9c2 cells transfected with the pUNO1-spike construct, when compared to control cells. Densitometry analysis showed elevation of GRP78 expression in spike gene-transfected cells, while housekeeping protein levels (GAPDH and Beta tubulin) remained constant, confirming equal protein loading (Fig. 8; Supplementary figure 8). The observed increase in GRP78 protein expression was consistent with the corresponding gene expression data.

**Figure 8:**
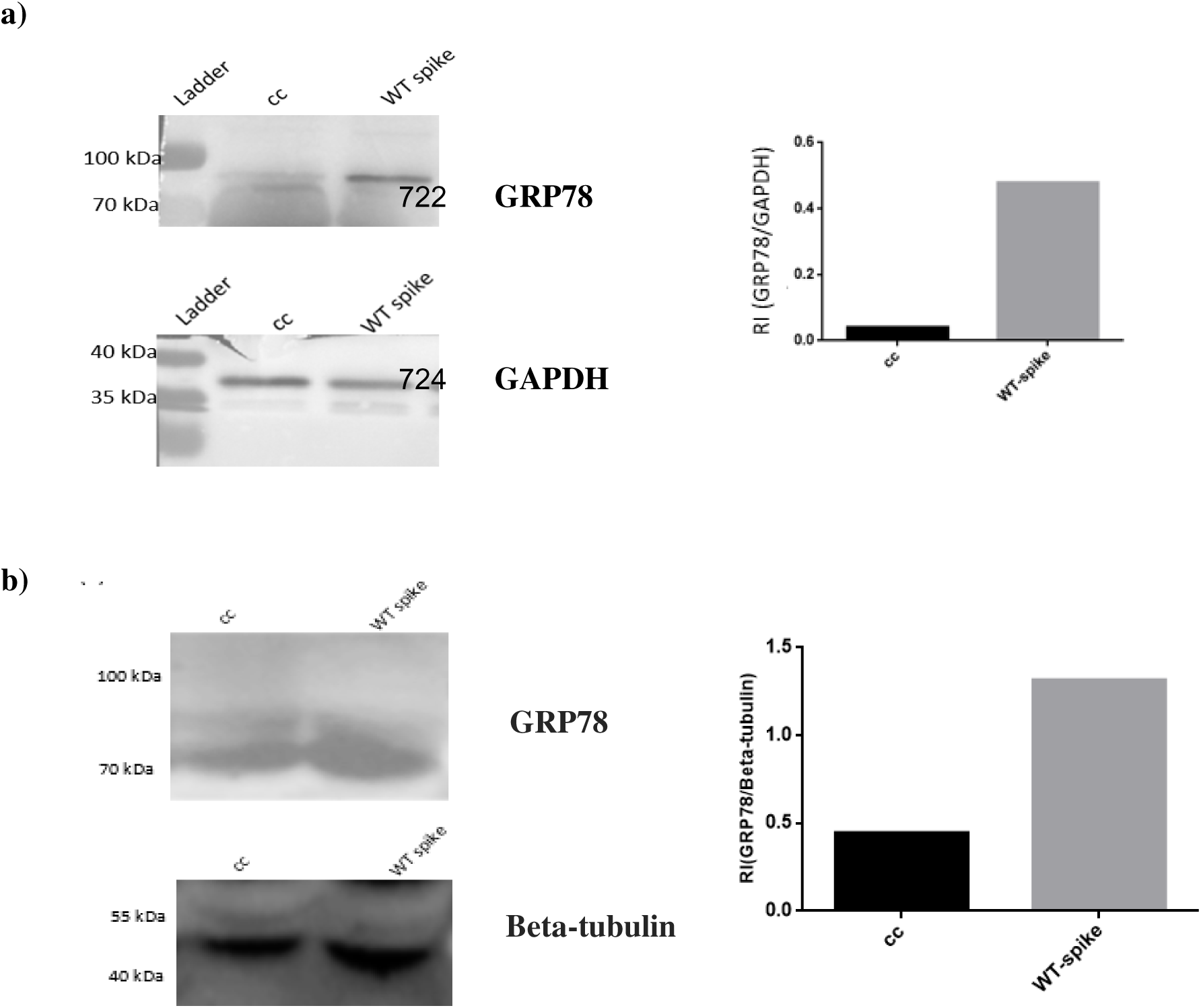
Immunoblot analysis of GRP78 in spike gene-transfected and control A549 and H9c2 cells. Densitometry quantification indicated that GRP78 (∼78 kDa) expression was elevated in spike-expressing (a) A549 cells and (b) H9c2 cell compared to controls, while GAPDH (36 kDa) and Beta-tubulin (‘50 kDa) levels remained consistent. CC=Cell control, WT=Wild type; RI=Relative intensity

#### GRP78 levels were elevated in serum samples from COVID-19 cases compared to those from Dengue cases and healthy individuals

COVID-19 (C12, C43, C22, C42, C4), Dengue (D2, D12, D3, D7, D9), and apparently healthy serum samples (H9 and CHS) were tested for GRP78 expression. GRP78 levels were markedly elevated in the COVID-19 serum samples. Quantitative assessment revealed an approximate two-fold increase in GRP78 expression in these samples compared to those obtained from Dengue cases and healthy controls (Fig. 9) (Ponceau S staining was used as a total protein loading control prior to immunodetection; Supplementary figure 6). These findings revealed that SARS-CoV-2 infection induced a more profound ER stress response compared to Dengue virus infection or in normal physiological conditions resulting in GRP78 overexpression and subsequent release into blood.

**Figure 9:**
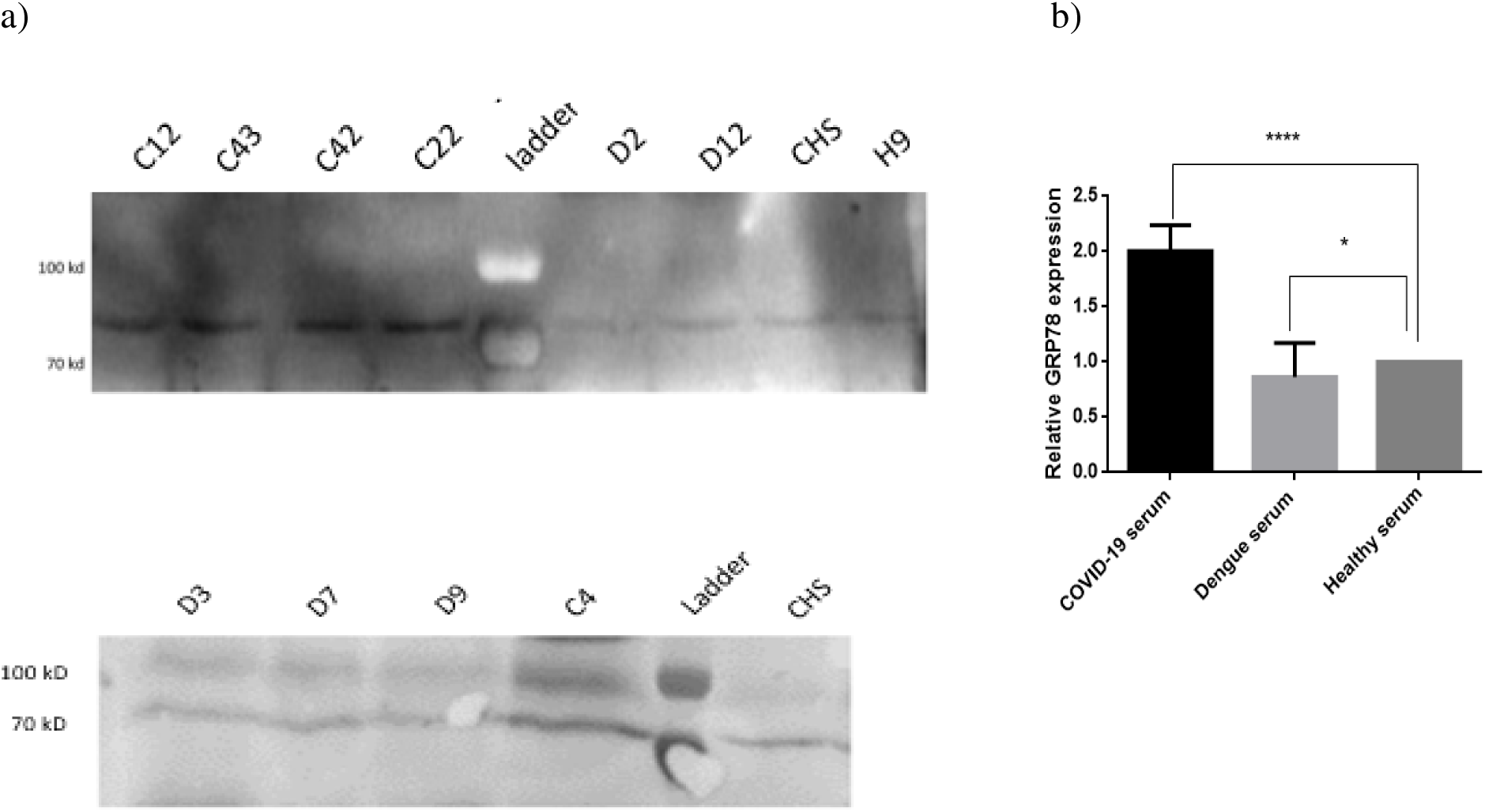
(a) Immunoblot analysis was performed on serum samples from COVID-19 cases (C12, C43, C42, C22, C4), Dengue cases (D2, D12, D3, D7, D9), and healthy controls (CHS, H9) using an anti-GRP78 antibody (b) Immunoblot analysis of GRP78 expression in serum samples from COVID-19 cases, Dengue cases (n = 5), and healthy controls using an anti-GRP78 antibody. (b) GRP78 band intensities were quantified by densitometry and expressed as relative expression normalized to the mean of healthy serum samples (set to 1). Bars represent mean ± SEM with individual serum samples shown. Statistical analysis was performed using a two-tailed one-sample *t* test comparing each group to the healthy reference value. GRP78 expression was significantly increased in COVID-19 serum compared with healthy controls (****p < 0.0001), whereas Dengue serum samples showed a moderate but significant reduction in GRP78 expression compared with healthy serum (*p* < 0.05).

#### Elevated GRP78 Levels in MI Serum Samples

The GRP78 expression significantly increased in MI serum samples compared with healthy controls. After normalization to the mean of healthy controls, the fold change value of GRP78 in individual MI affected individual’s serum clearly showed an increase, with most samples above the control baseline (Fig 10; Supplementary figure 5). The scattered distribution pattern showed variability among the individuals, but collectively, there was a visible increase in GRP78 in MI serum samples compared with normal controls, as depicted in the higher average level of its expression.

**Figure 10:**
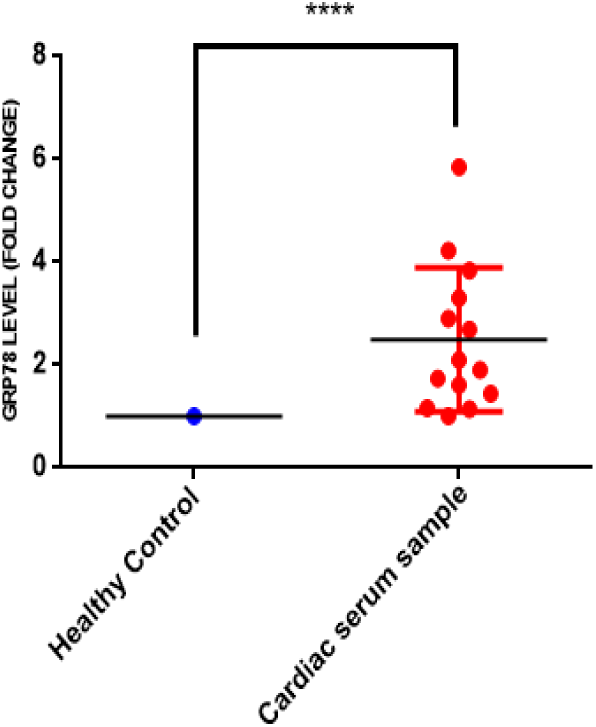
Increased GRP78 level in cardiac samples relative to healthy controls. Scatter dot plot showing individual fold-change values of cardiac serum samples normalized to the mean of healthy controls. Healthy control values represent the group average used for normalization, while each dot corresponds to an individual cardiac serum sample (n = 14). Horizontal bars indicate the mean ± 95% confidence interval. Cardiac serum samples showed a significant increase compared with healthy controls, as assessed by a one-sample t-test against a normalized value of 1.0 (t = 3.976, df = 13, p = 0.0016) and confirmed by a Wilcoxon signed-rank test (p = 0.0001). Statistical significance is indicated as **** (*p* < 0.0001).

## Discussion

The specific relationship between COVID-19 and heart-related anomalies appears to be multi-pronged. Although COVID-19 is primarily a respiratory sickness, many COVID-19 cases developed a new-onset of cardiac complications, even heart failure, during or after the illness.

C1q is involved in coagulation and production of atherosclerotic plaques^51^. In the present study, the average C1q level was found higher in COVID-19 compared to Dengue cases. Additionally, serum samples from COVID-19 or Dengue affected individuals had higher levels of C1q than those from apparently healthy individuals (Fig 1).

Circulating ICs (CICs) are formed when an Ab binds to an antigen. A strong correlation had been observed between CIC levels and complement activation^52^. Although the generation of ICs is an essential component of the immunological defense mechanism, excessive IC formation can be harmful to the host by interfering with normal physiological processes^53,54^. Our research findings revealed that COVID-19 and Dengue cases both exhibited C1C-C1q formation in 25 % and 42% affected individuals respectively, and in 32% of severe cases of COVID-19. These values were higher than the background CIC levels i.e. <10% of apparently healthy serums.

The SARS-CoV-2 spike protein binds strongly to human ACE2 and downregulates its expression^55^. ACE2 is part of the renin-angiotensin system (RAS), which includes angiotensinogen (AGT), Ang I, Ang II, renin, and ACE. Renin cleaves AGT to form inactive Ang I, which is converted by ACE into Ang II^56^. ACE2 (805 amino acids), a transmembrane protein, degrades Ang I to Ang-(1–9) and Ang II to Ang-(1–7). The latter is involved in vasodilation and vascular protection.

Reduced ACE2 activity leads to Ang II accumulation, promoting inflammation, fibrosis, vasoconstriction, and oxidative stress^57^. Ang II also enhances angiotensinogen expression via AT1R, creating a positive feedback loop^58,59^. Vasoconstriction will therefore, occur if Ang II level increases, which happens in case of COVID-19; this can provide a positive feedback loop for the production of more angiotensinogen, which, in turn, will make more Ang II.

Studies suggested a strong association between elevated AGT levels and increased COVID-19 disease severity^60^.

Our mass-spectrometry analysis showed that angiotensinogen expression in the COVID-19 serum samples was higher compared to the Dengue and healthy serum samples. The same was observed in another proteomics study on hospitalized COVID-19 serum samples^60^. They observed that AGT concentrations increased progressively from six severity groups from mild to critical cases. The rise in ANG was statistically significant, suggesting a strong correlation between AGT elevation and disease severity. These results pointed to RAS dysregulation as a possible contributor to COVID-19 pathophysiology, especially in severe cases, as discussed before.

A significant number of large population-based cohort studies had extensively documented the incidence of long-term health issues following SARS-CoV-2 infection, including cardiac complications, neuropsychiatric symptoms, autoimmune disorders, and chronic complement activation^61,62,63^. Similar long-term effects had also been observed in other acute infections, such as Dengue. Acute Dengue infection had also been shown to result in notable health consequences, with increased risks of neurological issues in the weeks following infection (7–14 days)^64,65^. However, the mechanisms through which these acute manifestations translated into chronic sequelae, particularly at the proteomic level remained unclear.

From disease-gene association enrichment analysis on the proteomics data, we observed that COVID-19 cases had a higher chance of developing thrombotic emboli, which could increase their vulnerability to diseases such as aortic valve disease or peripheral artery disease. Complement component 5 (C5) had been related to both coagulation system activation and platelet aggregation^66,67^. According to our proteomic data analysis, C5 in COVID-19 serums was significantly higher than in healthy serums. Our data also demonstrated that the anti-thrombin was down-regulated and coagulation factor XII was upregulated in COVID-19 compared to the Dengue, suggesting higher tendency of clot formation and thrombosis in COVID-19 cases.

In this context, it is noteworthy that apolipoprotein A1, which plays key roles in reverse cholesterol transport, cellular cholesterol homeostasis, and possesses anti-clotting and anti-aggregation properties, was decreased in COVID-19 serums^68^, but up-regulated during Dengue infections^69^. The lysis of clots was postponed by neutrophils and extracellular traps (NETs), and the viscous density of clots was enhanced in the vicinity of NETs^70^ Neutrophils promoted clotting and were usually higher in COVID-19 cases.

Increased level of C1q (as observed in our study) also bound to the fibrin helping in the clotting process. Circulating ICs and neutrophils may become embedded within vasoconstricted coronary and cerebral arteries. Once lodged, these complexes could be captured by coagulation factors such as C1q-bound fibrinogen, coagulation factor XII and other proteins involved in the coagulation cascade, thereby heightening the risk of myocardial infarction or cerebral thrombosis in COVID-19 cases (Fig. 10).

Circulating ICs had been also observed in Dengue in our study. This aligned with previous findings and supported our results^71^. Dengue infection results in vascular changes, such as vasodilation and increased permeability, which lead to hemorrhagic manifestations and plasma extravasation into the tissues. These features are characteristics of severe disease but can also occur at lower levels in mild and moderate disease states^72^. Our mass-spectrometry results supported the previous observations and showed that Dengue cases were more prone to bleeding, elevating their risk of developing anemia.

Angiotensinogen level was shown to be considerably lower in the case of Dengue^73^. Additionally, there was no downregulation of ACE2 expression, which maintained the balance between vasoconstriction (Ang II) and vasodilation (Ang-(1-7)). So, it is less likely that CICs will be able to accumulate or clog blood vessels which are already dilated and under hypotension in case of dengue infection.

In contrast to COVID-19, Dengue is associated with prolonged activated partial thromboplastin time (APTT), reduced fibrinogen levels, and elevated fibrinogen degradation products, indicating both impaired coagulation and enhanced fibrinolysis^74^.

On the other hand, COVID-19 cases showed markedly elevated fibrinogen levels, contributing to clot formation and hypo-fibrinolysis^75^. As mentioned before, even though CIC were observed in Dengue cases, the absence of vasoconstriction and the reduced levels of essential coagulation factors would further hinder their deposition and the development of thrombi in the coronary arteries.

Numerous studies had documented the pathogenic effects of spike protein itself. According to one research, the spike protein could damage the endothelium as observed in an animal model^76^. This toxicity applied to both severe forms and long COVID-19 as well as potentially to all spike vaccinations, based on the uncontrollably produced spike protein in the human system. Following mRNA vaccine injection, spike protein had been found to be present in significant amounts in free form circulating in blood, reaching various organs such as the liver, kidneys, brain, heart and lungs, in addition to being found on the cell surface^77^.

Therefore, we examined the impact of SARS-CoV-2 spike protein in three separate cell lines (lung, heart, and kidney cell lines) and found that the transfected spike protein was visible in these cell lines using confocal microscopy. Compared to the other two cell lines, the H9c2 (myoblast) exhibited a more concentrated deposition of spike protein in the cell membrane region.

To check for spike protein expression, Western blot was also performed from cell lysate and probed with S1 spike antibody. In HEK293T kidney cells, transfected spike protein S1 band was found in the appropriate region (100-130 kDa). However, we detected a band in the 70–100 kDa range in case of the other two cell lines, and upon performing in-gel digestion and mass spectrometry, we concluded that the band was most likely that of GRP78.

GRP78 is a stress-inducible chaperone. Evidence suggested that GRP78 could bind directly to the RBD of SARS-CoV-2 spike. Additionally, prior research had indicated that GRP78 acted as a crucial host auxiliary factor for the entry and infection of SARS-CoV-2^78^. SARS-CoV-2 replication was inhibited by knocking down GRP78 both *in vitro* and *in vivo*^79^.

Furthermore, higher plasma concentrations of GRP78/BiP were found in individuals with both subclinical atherosclerosis and metabolic problems; as a result, GRP78 was proposed as a relevant marker of cardiac and metabolic risk^80^. Our study showed a significant increase in GRP78 levels in MI serum samples compared with healthy controls (Supplementary figure 5), as evidenced by elevated normalized fold-change values and robust statistical significance.

We provided evidence that spike expression upregulated GRP78 gene in lung cells and protein expression in cardiac cells and lung cells. Interestingly, we observed that S1 Abs could also bind to GRP78 and could potentially form CICs. It had been reported that anti-GRP78 auto-Abs could induce endothelial cell activation and accelerate the development of atherosclerotic lesions^81^. So, it may be argued, that S1 Abs were analogous to anti-GRP78 Abs as both could bind to GRP78. Therefore, both could augment atherosclerosis which supported the clinical findings of coagulopathies in COVID-19 cases.

Finally, we observed significantly higher GRP78 expression directly in COVID-19 serum samples compared to those from Dengue cases and healthy controls, indicating that SARS-CoV-2 infection could induce a more pronounced endoplasmic reticulum (ER) stress response than Dengue virus infection or normal physiological conditions. Furthermore, our results showed that GRP78 auto-Abs and spike-abs could form ICs by directly binding to GRP78, abundantly present in blood and expressed by heart and lung tissues.

In conclusion, we have proposed two possible mechanisms that might be contributing towards the underlying pathophysiology of the cardiovascular manifestations of Long COVID. One mechanism involved circulating IC-associated emboli (like GRP78-spike Ab ICs in blood and tissues) which became trapped in already vasoconstricted blood vessels (due to reduced ACE2 expression and Ang II accumulation) and recruited complement components and other clotting-related factors; the entrapped ICs possibly elicited a type III hypersensitivity-like reaction, causing inflammatory and ischemic damages to the endothelial and cardiac tissues.

The other mechanism involved SARS-CoV-2 spike-mediated up-regulation of GRP78. The latter had been identified as additional receptor for SARS-CoV-2, besides ACE2. Over-expression possibly triggered formation of GRP78 auto-Abs. Furthermore, our results showed that spike Abs could cross-bind to GRP78 and therefore, might behave similar to GRP78 auto-Abs. These Ag-Ab complexes might bind to cell surface GRP78 in various tissues and organs and caused further tissue and blood vessel damages (type II hypersensitivity-like reactions).

Based on the information provided above, it appears that the production of immunological complexes occurs in both COVID-19 and Dengue. However, vasodilation happens in the case of Dengue resulting in generally observed hypotension (lower BP)^82,83^. Perhaps this renders it difficult for the ICs to become lodged in the dilated blood vessels.

**Figure 10:**
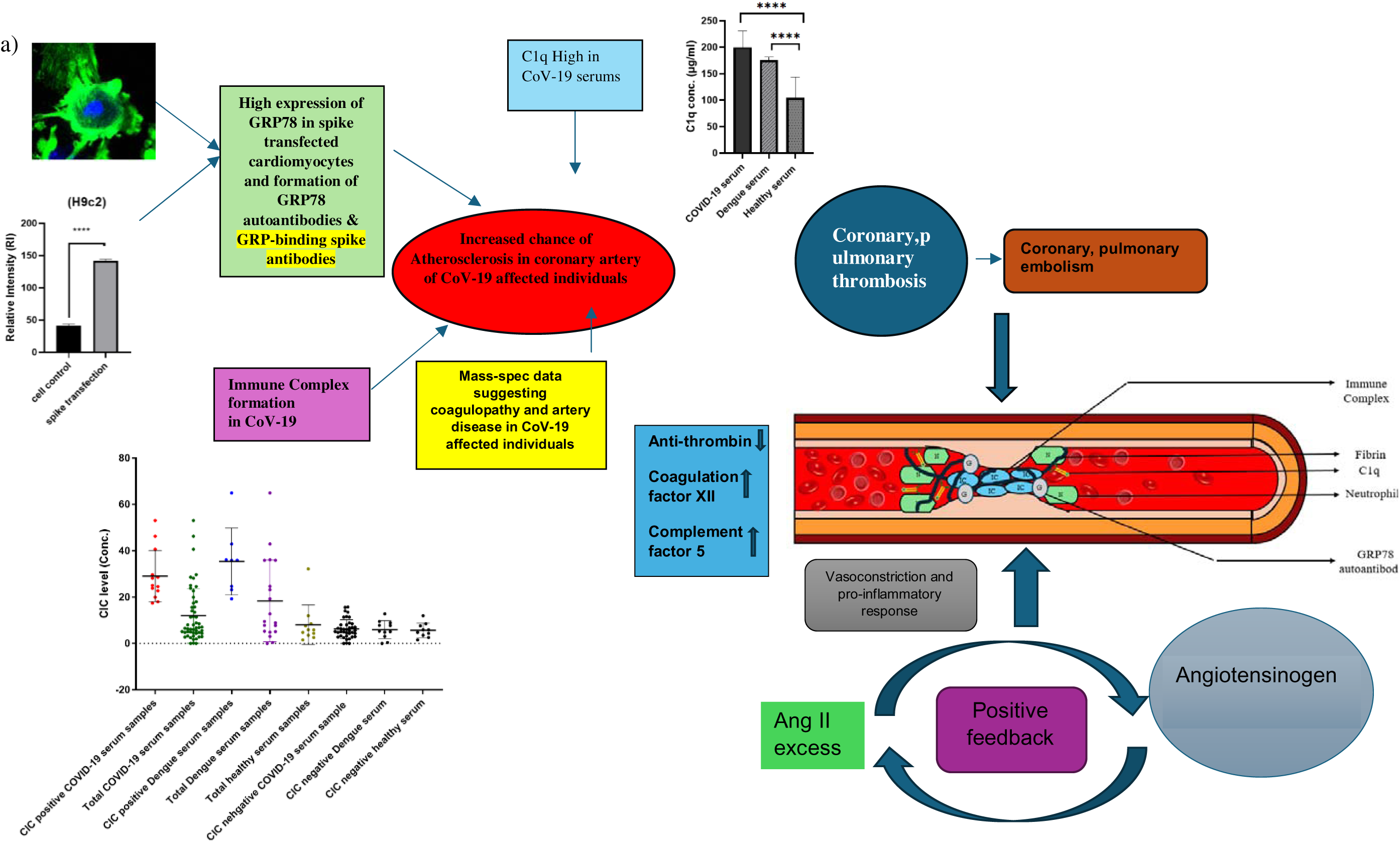
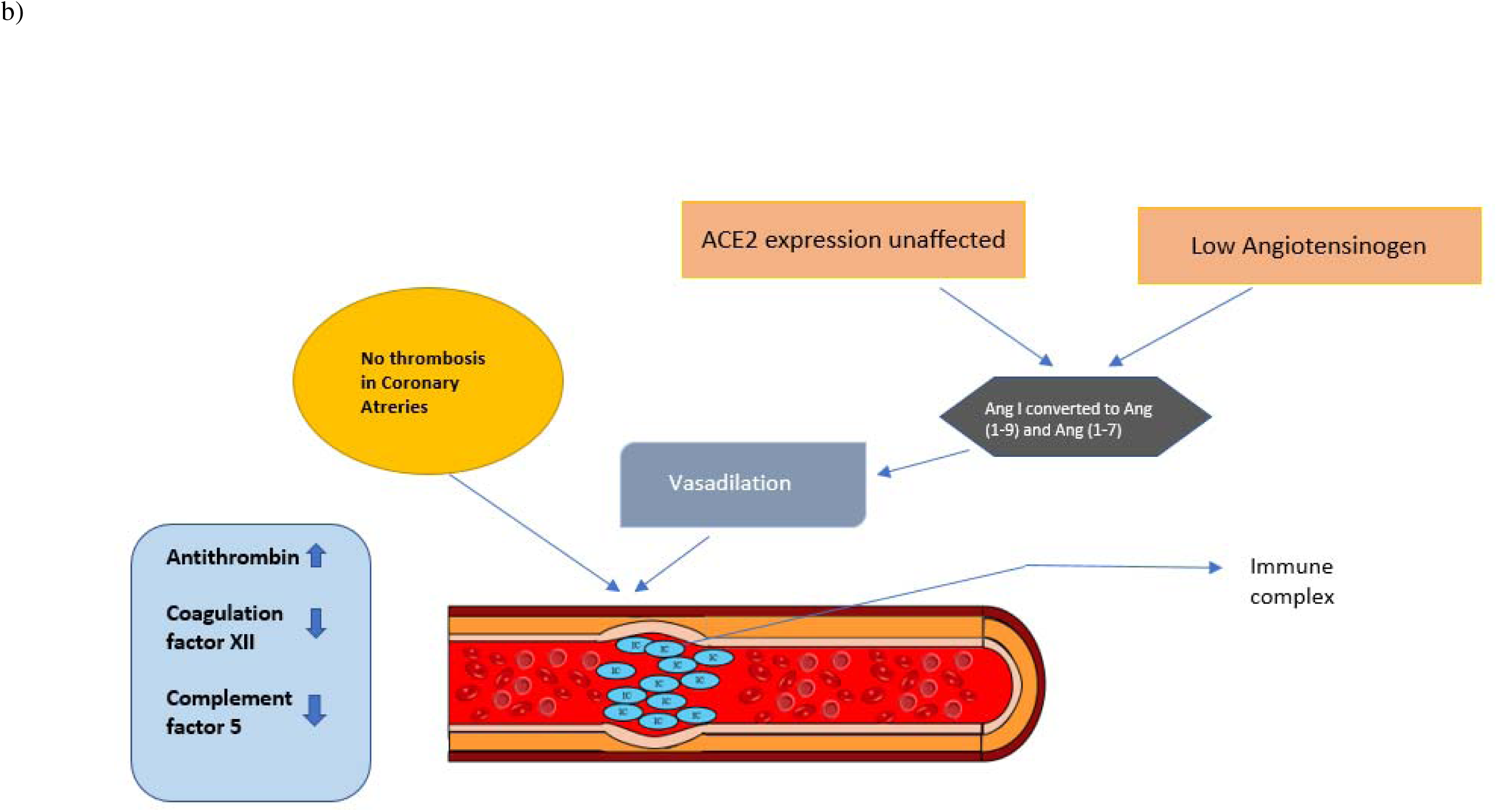
a) **Development of immune complexes and clotting in COVID-19 affected individuals’ coronary arteries.** Vasoconstriction and a pro-inflammatory response could arise from the failure to convert Ang II to Ang (1–7). Thus, an excess of Ang II may give positive feedback on the generation of Angiotensinogen, ultimately leading to vasoconstriction. Circulated immune complex and neutrophils can lodge in these vasoconstricted arteries. These lodged complexes can be trapped by coagulation factors, which include C1q-bound fibrinogen, coagulation factor-XII and other coagulation cascade-related proteins in coronary arteries, increasing the risk of heart attack in COVID-19 affected individuals. Additionally, SARS-CoV-2 spike protein has been implicated in the upregulation of glucose-regulated protein 78 (GRP78), which may enhance viral entry and stress responses. The immune response to the spike protein, including the generation of anti-spike antibodies and GRP78 autoantibodies, can lead to the formation of pathogenic antigen-antibody complexes with GRP78. These immune complexes may contribute to further vascular injury, inflammation, and thrombotic events, culminating in myocardial infarction and potentially cardiac arrest. b) **Immune Complex Formation Without Clotting in the Coronary Arteries of Dengue Affected individuals.** In Dengue affected individuals, circulating immune complexes can develop; however, unlike in COVID-19, these do not lead to significant clot formation in the coronary arteries. One contributing factor is the markedly lower levels of angiotensinogen observed in Dengue cases. Moreover, ACE2 expression remains unaffected or is not downregulated, which favors the conversion of angiotensin I (Ang I, 1–10) to vasodilatory peptides such as angiotensin (1–9) and angiotensin (1–7), rather than angiotensin (1–3). This biochemical environment promotes vasodilation rather than vasoconstriction. Additionally, Dengue is associated with a reduction in fibrinogen and coagulation factor XII levels, along with elevated antithrombin concentrations. These changes collectively result in a hypocoagulable state. As a consequence, despite the presence of circulating immune complexes, the lack of vasoconstriction and the depletion of key coagulation components prevent their deposition and the formation of thrombi within the coronary arteries of Dengue affected individuals.

Vasoconstriction occurs in conjunction with clotting factor elevation in case of COVID-19. Individuals with COVID-19 might have circulating ICs that are stuck in these constricted arteries, increasing their risk of heart attacks and other cardiac problems. Although BP data were not available for the COVID-19 affected individuals in our study, a recent report showed that COVID-19 affected individuals (irrespective of severity) consistently showed higher BP, arterial stiffness and accelerated vascular ageing compared to controls up to 6-12 months after COVID-19^84^. Even though COVID-19 had been largely a respiratory illness, a considerable proportion of individuals experienced heart failure that developed during the illness. Therefore, it is critical to comprehend the underlying mechanisms and cardiac difficulties induced by SARS-CoV-2.

## Supporting information

Similarly, archived serum samples from 19 Pre COVID-19 pandemic Dengue cases were collected from Apollo Multispecialty Hospital in 2017

## Acknowledgements

This study was funded by CSIR- India (grant numbers: MLP 130 and OLP 118) (CSIR Digital Surveillance Vertical for COVID- 19 mitigation in India). The grants were given to S.B. The funders had no role in the study design, in the collection, analysis and interpretation of data; in the writing of the manuscript; and in the decision to submit the manuscript for publication. The authors acknowledge the support received from the Director of CSIR- IICB. S.B. also acknowledges AcSIR for the support. S.S., T.R. and A.M. acknowledge the support of CSIR for their CSIR Senior Research Fellowship; S.D. acknowledges UGC for his Senior Research Fellowship. The authors would also like to acknowledge the use of relevant facilities at CSIR-IICB including the Central Instrumentation Facility (CIF).

## Author contributions

S.B. conceived and designed the study. A.B. critically reviewed the study. S.S. conducted the majority of the experiments. The proteomic work was carried out by T.R. Archived serum samples with necessary information were obtained by A.D., S.G. and S.D.T. The original draft of the manuscript was written by S.S., A.M., T.R., S.D. and S.B. Critical data analysis was performed by S.B. and S.S. Funding for the study was acquired by S.B. All authors reviewed and approved the final manuscript.

## Ethical statement

The study was conducted according to the guidelines of the Declaration of Helsinki and approved by the Institutional Ethical Committees of Apollo Multispecialty Hospital, Kolkata, Calcutta National Medical College and Hospital, Kolkata, and the Council of Scientific and Industrial Research- Indian Institute of Chemical Biology, Kolkata.

## Consent to publish

Written informed consent (in their native language) was obtained from all individual participants included in this study. All experiments were carried out in accordance with the relevant guidelines and regulations.

