## Supplementary material for "Spike antibodies targeting GRP78 predispose to cardiovascular complications compared to Dengue": Similarly, archived serum samples from 19 Pre COVID-19 pandemic Dengue cases were collected from Apollo Multispecialty Hospital in 2017

**Supplementary Information:**

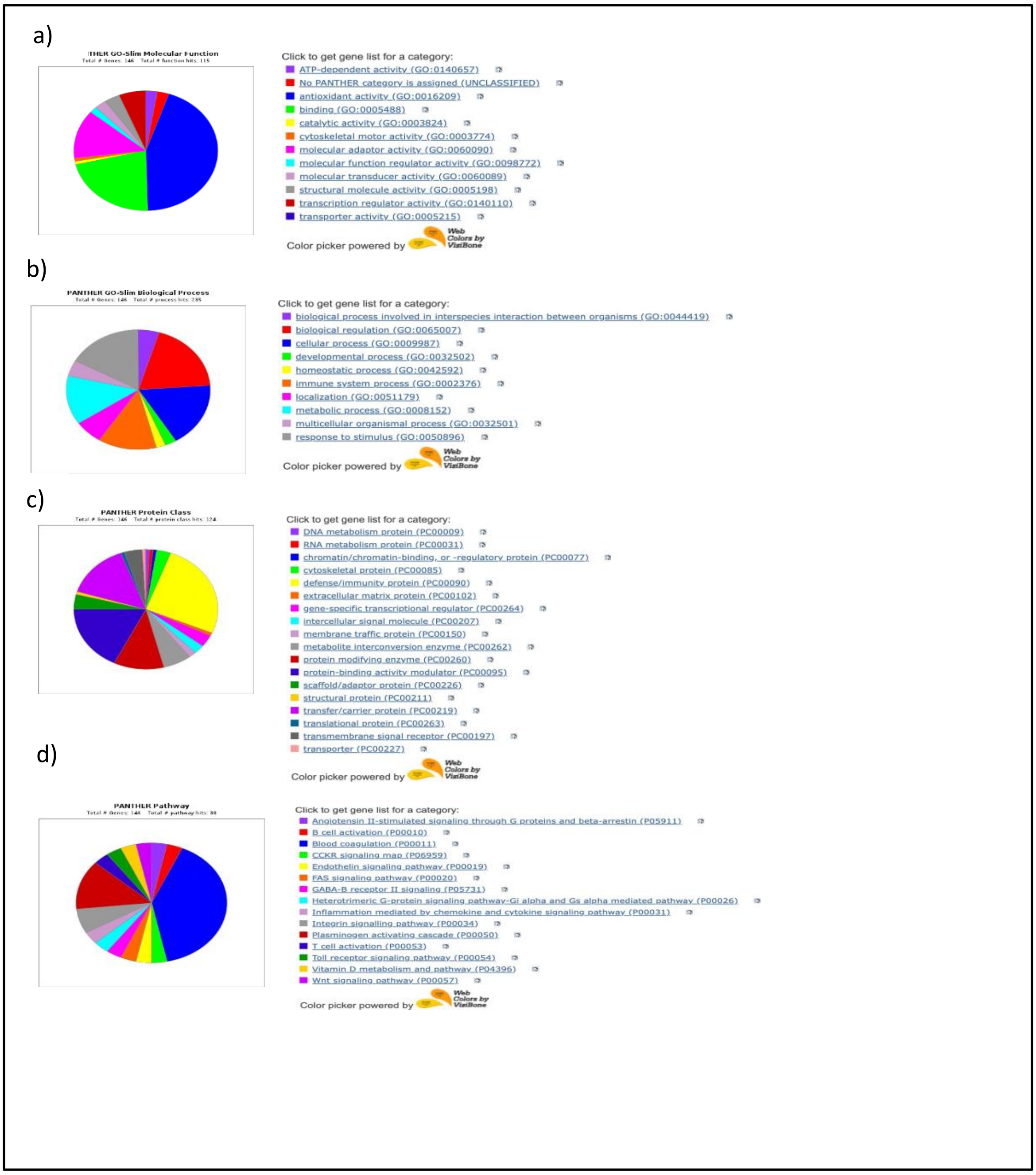

### Supplementary figures

**Supplementary figure 1: Disease-specific proteomic classification of COVID-19-associated proteins.** Functionally annotated proteins identified in COVID-19 samples were categorized using Gene Ontology (GO) and PANTHER classification systems. (a) **Molecular Function:** Classification of proteins based on their biochemical activity, including catalytic activity, binding, and receptor activity. (b) **Biological Process:** Distribution of proteins across major biological processes such as immune response, inflammation, and metabolism. (c) **PANTHER Protein Class:** Categorization of proteins into structural and functional classes, including enzymes, transporters, and signaling molecules. (d) **PANTHER Pathway:** Mapping of proteins to key signaling and metabolic pathways affected during SARS-CoV-2 infection. This multi-level annotation highlights the molecular complexity and diverse functional roles of proteins altered in COVID-19.

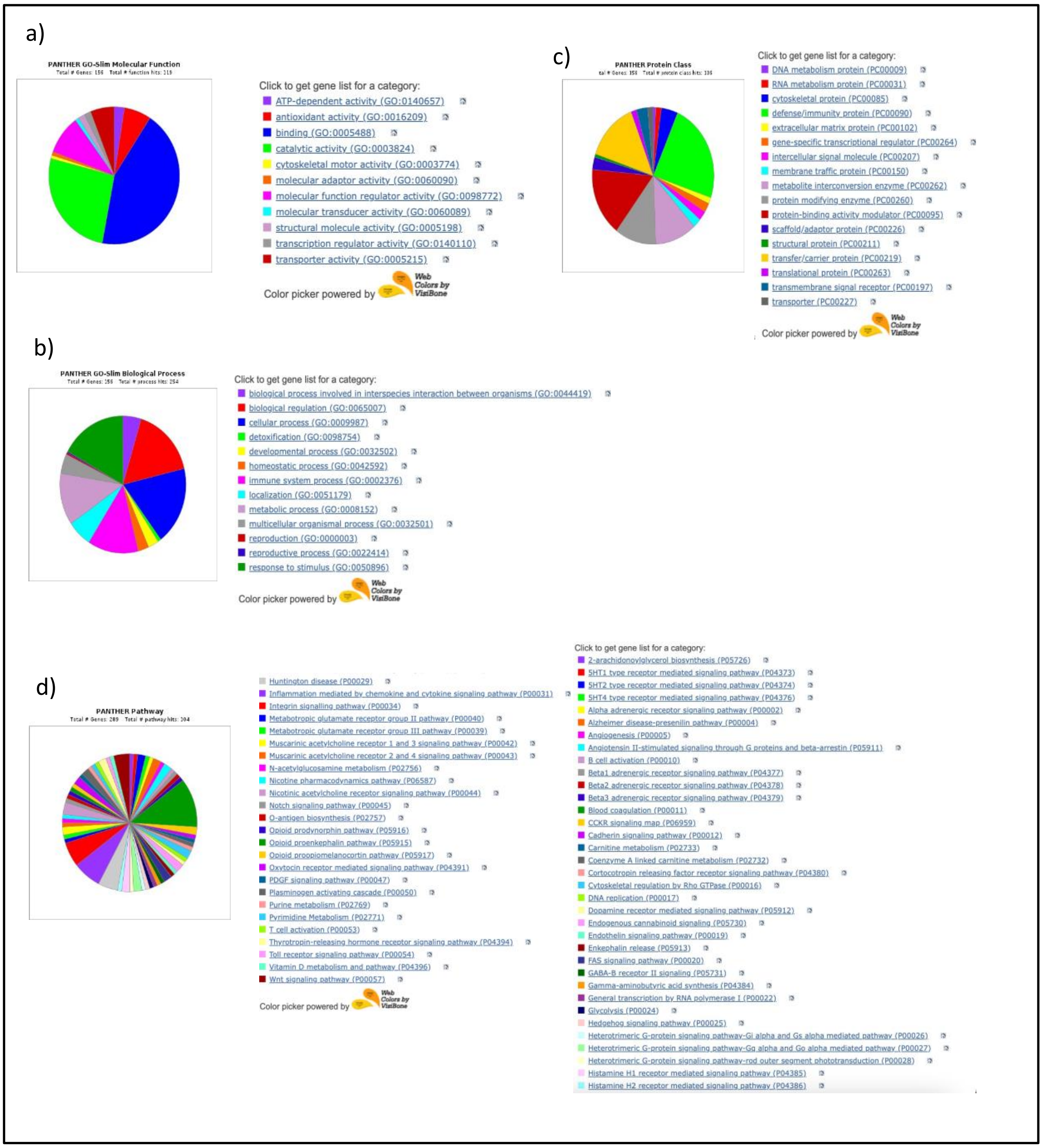

**Supplementary figure 2: Disease-specific proteomic classification of Dengue-associated proteins.** Functionally annotated proteins identified in Dengue were categorized using Gene Ontology (GO) and PANTHER classification systems. (a) **Molecular Function:** Classification of proteins based on their biochemical activity, including catalytic activity, binding, and receptor activity. (b) **Biological Process:** Distribution of proteins across major biological processes such as immune response, inflammation, and metabolism. (c) **PANTHER Protein Class:** Categorization of proteins into structural and functional classes, including enzymes, transporters, and signaling molecules. (d) **PANTHER Pathway:** Mapping of proteins to key signaling and metabolic pathways affected during Dengue infection. This multi-level annotation highlights the molecular complexity and diverse functional roles of proteins altered in Dengue.

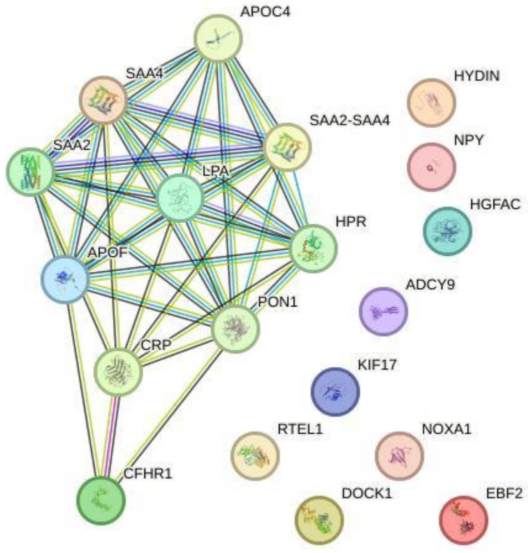

**Supplementary figure 3**: STRING-based network analysis of proteins uniquely dysregulated in COVID-19 infection. The network showed a highly significant PPI enrichment ($p = 1.11 \times 10^{-16}$), supporting the presence of functional connectivity and shared biological roles among these proteins in the context of COVID-19 pathogenesis.

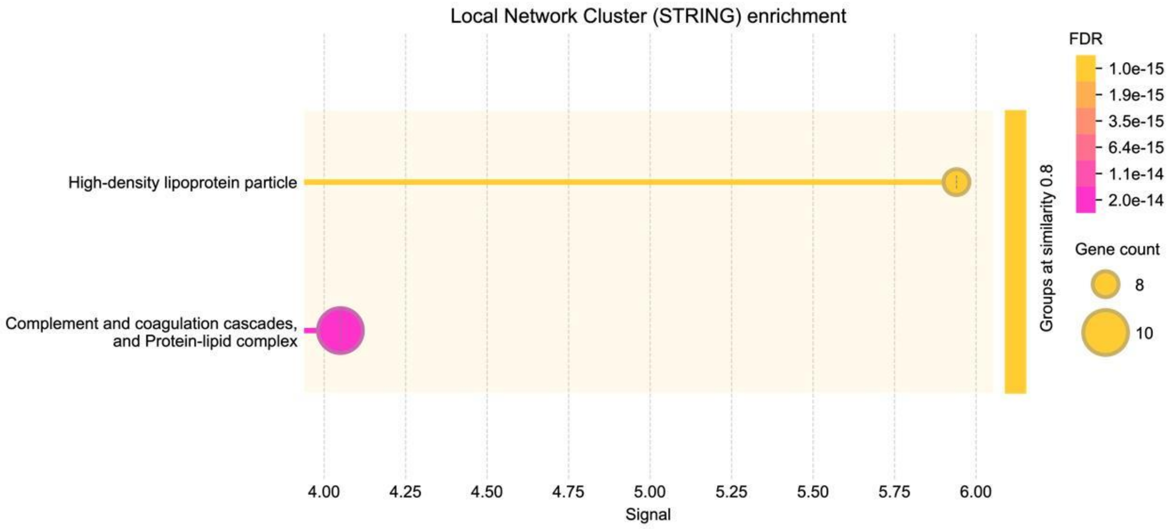

**Supplementary figure 4:** Sub-clustering of dysregulated proteins reveals distinct functional modules. The protein-protein interaction network of proteins uniquely altered in COVID-19 infection further resolves into two major local clusters. These clusters are enriched for proteins commonly associated with high-density lipoproteins (HDL), complement and coagulation cascades, and protein-lipid complexes.

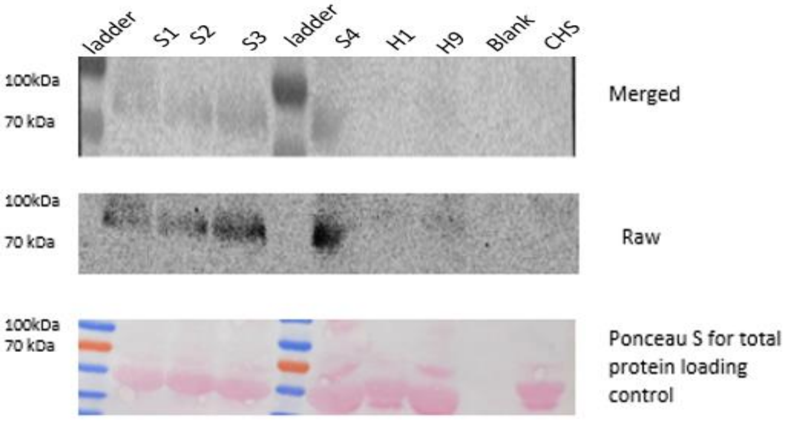

**Supplementary figure 5**:

Representative Immunoblot indicating GRP78 protein expression in MI serum samples and healthy controls. Serum samples from healthy controls (H1, H2 and CHS) and cardiac patients (S1, S2, S3 and S4) were resolved by SDS–PAGE and probed with anti-GRP78 antibody. The top panel shows the merged image, the middle panel depicts the raw GRP78 signal, and the bottom panel shows Ponceau S staining used as a total protein loading control. This representative blot indicates the increased GRP78 levels observed in MI serum samples relative to healthy controls.

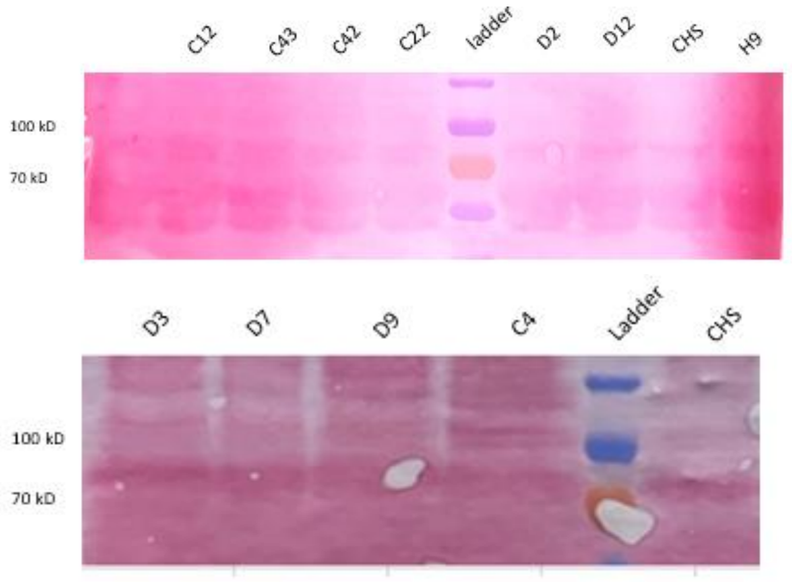

**Supplementary figure 6:** Ponceau S staining of the nitrocellulose membrane corresponding to the GRP78 immunoblots shown in Figure 9. The entire membrane is shown to demonstrate uniform protein transfer and comparable total protein loading across all serum samples, including COVID-19 cases (C12, C43, C42, C22), Dengue cases (D2, D12, D3, D7, D9), and healthy controls (CHS, H9). Ponceau S staining was used as a total protein loading control prior to immunodetection.

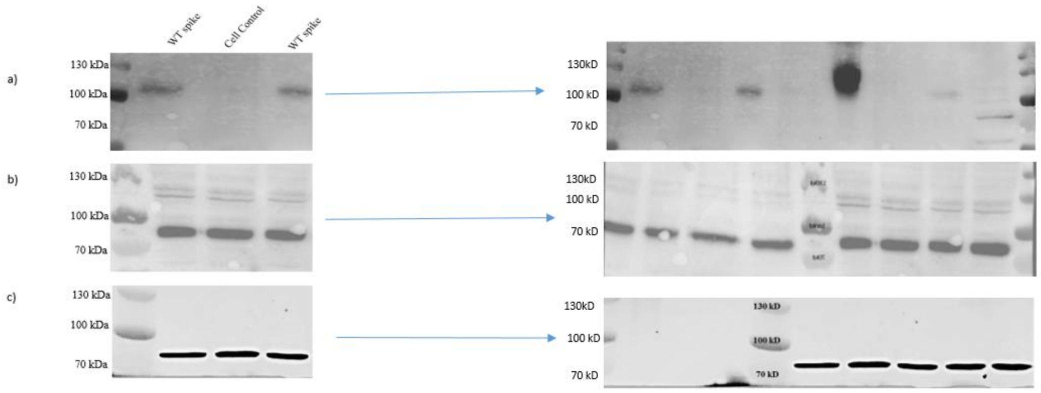

**Supplementary figure 7:** Full-length immunoblots corresponding to Figure 3, demonstrating detection of SARS-CoV-2 spike protein and cell control in (A) HEK293T, (B) H9c2, and (C) A549 cell lysates collected 72 hours post-transfection.

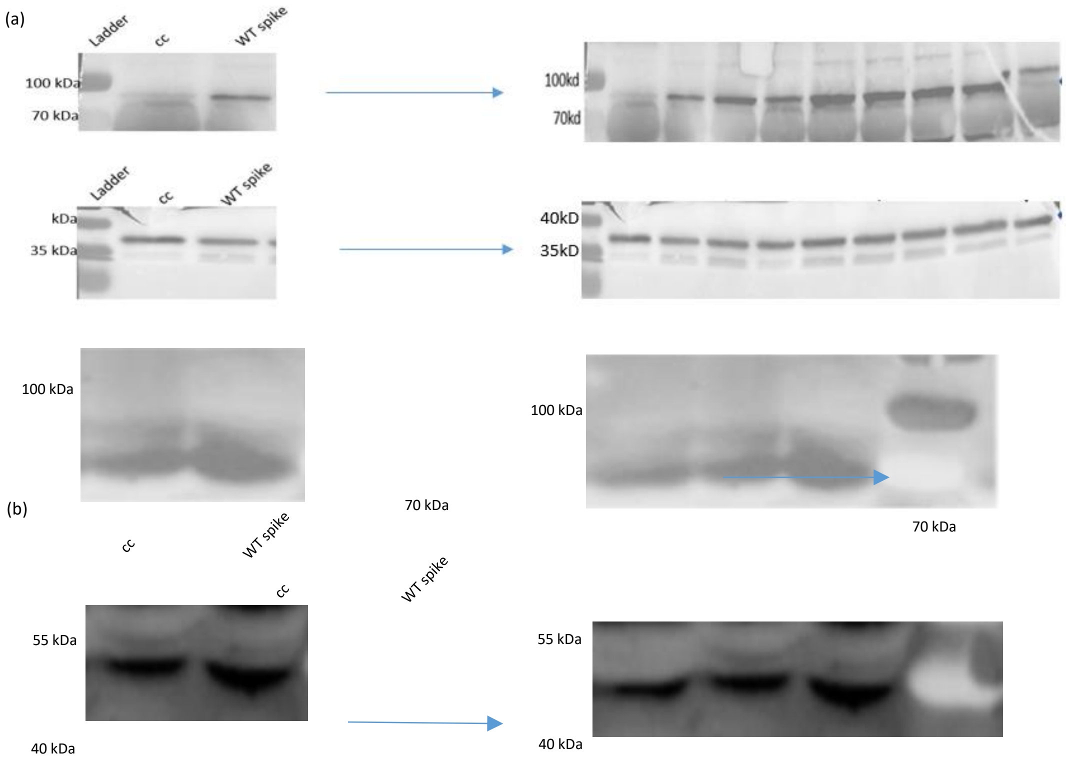

**Supplementary figure 8:** Full-length immunoblots corresponding to Figure 8, demonstrating spike transfected GRP78 expression in (a) A549 and (b) H9c2 cell line.

| Serial no. | MS no. | Sex | Ward status | NS1 antigen | DV IgM | DV IgG |
|---|---|---|---|---|---|---|
| 1 | D1 | M | STD | (+) | (+) | (+) |
| 2 | D2 | M | STD | (+) | (+) | (-) |
| 3 | D3 | F | STD | (+) | (-) | (+) |
| 4 | D4 | M | STD | (+) | (+) | (-) |
| 5 | D5 | F | STD | (+) | (-) | (+) |
| 6 | D6 | F | STD | (+) | (+) | (-) |
| 7 | D7 | F | STD | (+) | (+) | (-) |
| 8 | D8 | M | STD | (+) | (+) | (-) |
| 9 | D9 | M | STD | (+) | (-) | (+) |
| 10 | D10 | M | STD | (+) | (+) | (+) |
| 11 | D11 | M | STD | (+) | (+) | (+) |
| 12 | D12 | F | STD | (+) | (+) | (+) |
| 13 | D13 | M | STD | (+) | (+) | (+) |
| 14 | D14 | F | STD | (+) | (+) | (+) |
| 15 | D15 | M | STD | (+) | (-) | (+) |
| 16 | D16 | F | STD | (+) | (+) | (-) |
| 17 | D17 | M | STD | (+) | (-) | (+) |
| 18 | D18 | F | STD | (+) | (-) | (+) |
| 19 | D19 | F | STD | (+) | (-) | (+) |

**Supplementary table 1:** Serological profile of dengue patients showing NS1 antigen, dengue virus (DV) IgM, and DV IgG status. All patients were NS1 positive, indicating acute dengue infection.

| Serial no | MS no | Sex | Ward status | Vaccination history | **AbCheck IgM (15min)** | **AbCheck IgG (15 min)** | **AbCheck IgM (24 hr)** | **AbCheck IgG (24 hr)** |
|---|---|---|---|---|---|---|---|---|
| 1 | S1 | M | HDU | Yes | (-) | (+) VVF | (-) | (+) VVF |
| 2 | S2 | F | HDU | No | (-) | (+) | (-) | (+) |
| 3 | S3 | M | ICU | No | (-) | (-) | (+) VVF | (+) VVF |
| 4 | S4 | M | ICU | Yes (Covishield) | (-) | (+) | (+) VF | (+) |
| 5 | S5 | F | ICU | Yes (Covishield) | (-) | (-) | (-) | (-) |
| 6 | S6 | M | ICU | Yes (Covishield) | (+) VF | (-) | F (+) | (+) VF |
| 7 | S7 | F | HDU | Yes (Covishield) | (-) | (-) | (+) | (+) |
| 8 | S8 | F | ICU | Yes (Covishield) | (-) | (+) VF | (+) | (+) |
| 9 | S9 | M | HDU | Yes (Covaxin) | (+) VVF | (+) VF | F (+) | (+) |
| 10 | S10 | M | HDU | Yes (Covishield) | (-) | (-) | (-) | (+) VF |
| 11 | S11 | F | ICU | NA | (-) | (+) | F (+) | (+) |
| 12 | S12 | M | ICU | Yes (Covishield) | (-) | F (+) | (-) | F (+) |
| 13 | S13 | F | HDU | Yes (Covishield) | (-) | (-) | F (+) | F (+) |
| 14 | S14 | F | HDU | Yes (Covishield) | (-) | (-) | (-) | (F+) |

**Supplementary table 2:** List of serum samples collected from myocardial infarction (MI) patients, used in GRP78 western blot analysis

| Sample type | Sl no. | Manuscript name | Pride Proteome data base name | Sex | Severity |
|---|---|---|---|---|---|
| COVID-19 serum | 1 | C1 | 280B | F | STD |
| | 2 | C3 | 468_6 | M | STD |
| | 3 | C4 | TKN3 | F | STD |
| | 4 | C5 | TKN6 | F | STD |
| | 5 | C6 | TKN (merged) | F | STD |
| | 6 | C7 | | F | STD |
| | 7 | C8 | | M | STD |
| | 9 | C11 | | M | STD |
| | 10 | C12 | | M | STD |
| Dengue Serum | 1 | D1 | TKDV1 | M | STD |
| | 2 | D2 | TKDV4 | M | STD |
| | 3 | D3 | 107ZD | F | STD |
| | 4 | D4 | 138ZD | M | STD |
| | 5 | D5 | DEN (merged) | F | STD |
| | 6 | D6 | | F | STD |
| | 7 | D7 | | F | STD |
| | 8 | D8 | | M | STD |
| | 9 | D9 | | M | STD |
| | 10 | D10 | | M | STD |
| Healthy serum | 1 | H1 | TKN4 | F | STD |
| | 2 | H2 | TKN8 | F | STD |
| | 3 | H3 | 119ZD | M | STD |
| | 4 | H4 | Normal | M | STD |
| | 5 | H5 | | F | STD |
| | 6 | H6 | | M | STD |
| | 7 | H7 | | M | STD |
| | 8 | H8 | | F | STD |
| | 9 | H9 | | M | STD |

**Supplementary table 3:** Annotation of serum samples used in the study, showing manuscript identifiers and corresponding PRIDE Proteome database names. COVID-19, Dengue, and Healthy serum samples are listed along with donor sex and disease severity (STD). **Both individual serum samples and merged (pooled) serum samples were used for analysis.** Samples indicated as merged represent pooled sera from multiple individuals, whereas all other entries correspond to individually processed serum samples.
